# ecDNA-driven oncogene super-expressors shape immunoevasive tumor microenvironment

**DOI:** 10.1101/2025.11.15.688565

**Authors:** Kailiang Qiao, Qing-Lin Yang, Tuo Li, Xiongfeng Chen, Zeynep Yazgan, Yoon Jung Kim, Collin Gilbreath, Jun Yi Stanley Lim, Yipeng Xie, Xiaohui Sun, Yang Liu, Yiyue Jia, Zhijian J. Chen, Huocong Huang, Sihan Wu

## Abstract

ecDNA contributes to cancer genetic heterogeneity through random segregation during mitosis. Emerging evidence links ecDNA to immune evasion, but the mechanism remains elusive. Using genetically engineered mouse models of pancreatic ductal adenocarcinoma (PDAC), we show that *Kras* and *Myc* oncogenes are amplified either on ecDNAs or as homogeneously staining regions (HSRs) on chromosomes. ecDNA-driven tumors are more aggressive in immunocompetent mice. Single-cell transcriptomic and histological analyses reveal that ecDNA-driven tumors rapidly establish an immunoevasive tumor microenvironment (TME), marked by increased myofibroblastic cancer-associated fibroblasts (myCAFs) and reduced T cell infiltration. Mechanistically, ecDNA heterogeneity generates a subset of cancer cells with extremely high *Kras* expression, termed super-expressors, which secrete amphiregulin to promote myCAF expansion and suppress T cell infiltration. Clonally organized super-expressors establish an immunoevasive niche in the TME from patients with PDAC. Our findings demonstrate a causal role of ecDNA in TME remodeling, offering insights into cancer heterogeneity and immune evasion.

## Introduction

Cancers possess multiple mechanisms to evade immune surveillance and destruction. Cancer cell-intrinsic alterations, such as genetic, epigenetic, and metabolic aberrations, profoundly influence the composition and function of the tumor microenvironment (TME).^1,2^ For example, many canonical oncogenes, such as *KRAS* and *MYC*, have been shown to facilitate immune evasion by regulating the expression of cytokines, chemokines, and immune checkpoint molecules.^3–7^ Consequently, the cell composition and cell-cell communication in the TME are reshaped to be more immunosuppressive, characterized by decreased T cell infiltration, increased expansion of cancer-associated fibroblasts (CAFs), and, eventually, reduced efficacy of immune therapies.

Extrachromosomal DNA (ecDNA) represents an emerging hallmark of aggressive cancer. Its non-Mendelian inheritance, high copy number, and hyper-accessible chromatin drive profound oncogene amplification and intratumoral heterogeneity.^8,9^ Patients with ecDNA-driven cancers usually have poor clinical outcomes, including therapy resistance and shorter survival.^10,11^

More recently, ecDNA amplification has been increasingly associated with immunoevasive TME. Two pioneering pan-cancer analyses using The Cancer Genome Atlas (TCGA) data, based on bulk-cell RNA sequencing (RNA-seq), independently demonstrate that ecDNA-containing tumors exhibit an immune-cold phenotype, characterized by reduced immune cell infiltration, downregulated immunoregulatory pathways, and diminished antigen presentation.^12,13^ Aligned with these findings, a single-cell RNA sequencing (scRNAseq) study reveals that urothelial carcinomas harboring ecDNA display decreased MHC-I expression in malignant cells and increased regulatory T cell (Treg) infiltration compared to tumors lacking focal somatic copy number amplifications.^14^ A spatial transcriptomic study in pancreatic ductal adenocarcinoma (PDAC) further links high MYC expression, likely through the amplification of ecDNA, to the enrichment of CD90+ myofibroblastic CAFs (myCAFs) and the depletion of cytotoxic T cells.^15^ Notably, ecDNA may also harbor immunomodulatory genes independent of canonical oncogenes, suggestive of positive selection of immunoevasive ecDNA species.^16–18^ Bulk-cell transcriptomic analysis shows that tumors carrying such immunomodulatory ecDNAs exhibit significantly reduced T cell infiltration compared to those harboring oncogene-only ecDNAs.^18^

While accumulating evidence links ecDNA to an immunoevasive phenotype, the causality underlying this association remains elusive: Does ecDNA shape an immunosuppressive TME, or does a more immunosuppressive TME favor ecDNA pathogenesis? Addressing this question faces significant challenges. First, clinical data relying on bulk-cell sequencing cannot accurately deconvolute the cellular complexity of the TME. Second, while emerging single-cell and spatial genomic technologies offer high-resolution insights, establishing causality in human tissues remains a formidable challenge. This difficulty stems from the lack of isogenic controls, which makes it extremely hard to disentangle the specific effects of ecDNA from other forms of chromosomal amplification. Finally, while mouse ecDNA cancer models become available,^19^ current studies have yet to generate isogenic controls comparing ecDNA to chromosomal amplification.

To address these limitations, we used the well-established *KPfC* mouse PDAC model (*Kras^LSL-G12D/+^*; *Trp53^fl/fl^*; *Pdx1^Cre/+^*) to generate primary isogenic cancer cell clones.^20–22^ Surprisingly, we found that *Kras* and *Myc* oncogenes were spontaneously amplified as ecDNAs and homogeneously staining regions (HSRs) on chromosomes. From clones typically harboring both forms of amplification, we isolated isogenic pairs carrying exclusively ecDNA or HSR. Using scRNAseq, genetic knockout, and histological analyses for tumors in immunocompetent syngeneic mice, together with spatial transcriptomic analyses in PDAC patient samples, we investigated the causal role of ecDNA amplification in shaping an immunoevasive TME.

## Results

### *KPfC* PDAC spontaneously acquires ecDNA and HSR amplification

To establish a comprehensive library of congenic cancer cell clones, we harvested fully developed tumors that had been raised spontaneously in the *KPfC* mouse and processed them into single-cell suspensions through enzymatic digestion. Individual clones were isolated through limiting dilution and subsequently expanded through serial passaging. Each established clone was rigorously characterized and validated for tumorigenic potential through subcutaneous injection into immunocompetent syngeneic recipients, confirming their ability to form tumors *in vivo*. This library of clones recapitulates the heterogeneity of cancer cell populations present within the entire tumor, capturing distinct lineage phenotypes that represent the diverse cellular subpopulations found in the spontaneous tumors. This approach yielded a genetically homogeneous yet phenotypically diverse collection of tumor cell clones that retained the key molecular characteristics of the parental tumors (Figure 1A).

**Figure 1.**
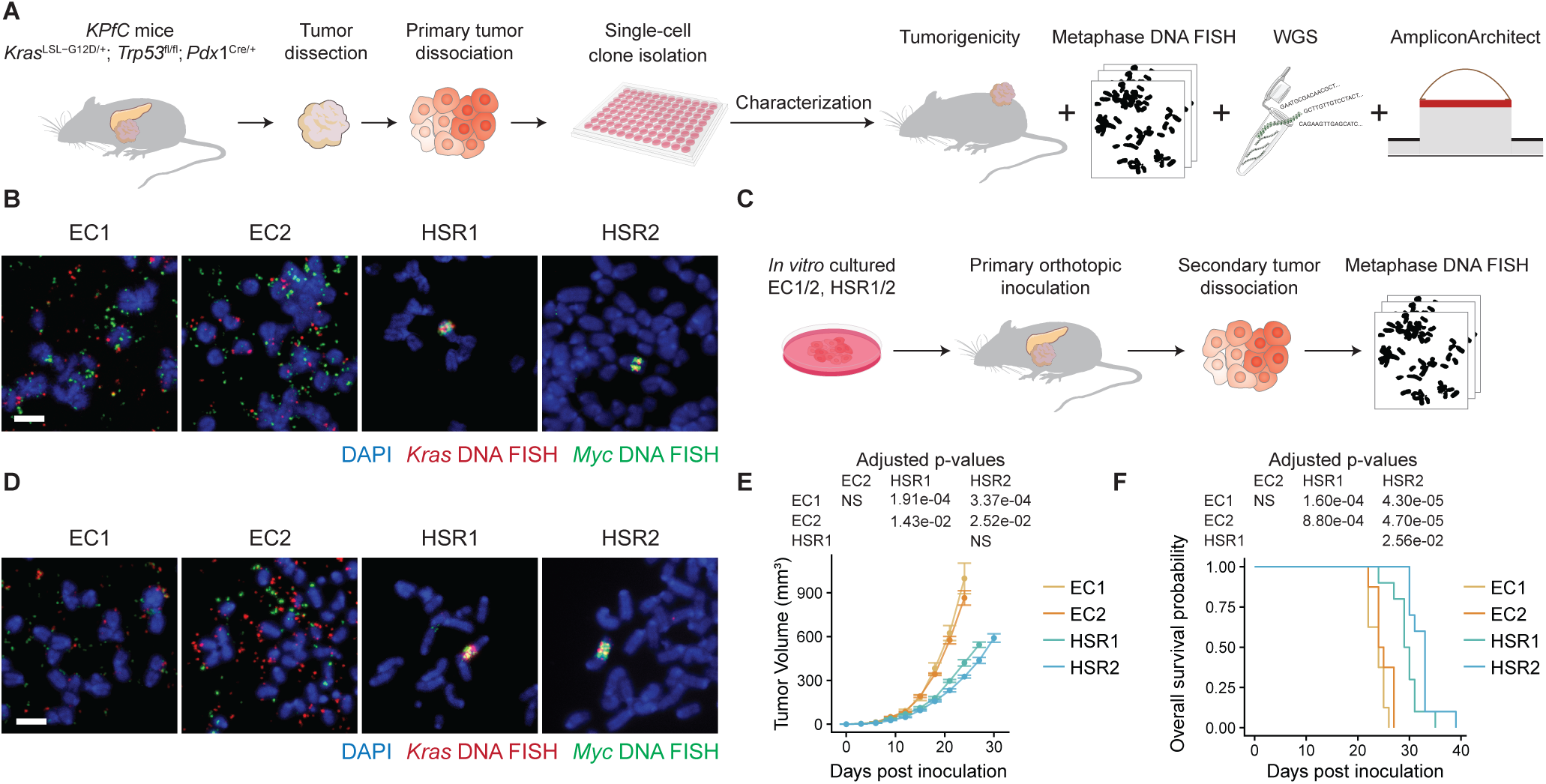
Establishment and characterization of ecDNA and HSR-containing PDAC cell lines. (A) Flowchart depicting the process to isolate and identify ecDNA or HSR-containing isogenic PDAC cell clones from tumors in *KPfC* mice. (B) *Kras* and *Myc* DNA FISH at metaphase spreads in EC1/2 and HSR1/2 *in vitro* cultured cells from the primary tumor dissociation. Scale bar: 5 μm. (C) Workflow for validating the maintenance of ecDNA and HSR amplicons in orthotopic tumors derived from EC1/2 and HSR1/2 clones. (D) *Kras* and *Myc* DNA FISH at metaphase spreads in EC1/2 and HSR1/2 *in vitro* cultured cells from the secondary tumor dissociation. Scale bar: 5 μm. (E) Subcutaneous tumor growth curve of EC1/2 and HSR1/2 isogenic clones. Two-way ANOVA with Tukey’s HSD. NS indicates not significant. Sample size: EC1, n = 9; EC2, n = 8; HSR1, n = 9; HSR2, n = 10. (F) Overall survival of C57BL/6J mice bearing EC1/2 or HSR1/2 tumors. Pairwise log-rank test with false discovery rate correction. Sample size: EC1, n = 8; EC2, n = 8; HSR1, n = 10; HSR2, n = 10.

We identified one clone, designated CT1BA5, that harbored both *Kras* ecDNAs and HSRs.^23^ We then isolated two ecDNA-only and two HSR-only isogenic subclones from the parental CT1BA5 cells, termed EC1 or EC2 and HSR1 or HSR2. All isogenic subclones were verified by *Kras* DNA fluorescence *in situ* hybridization (FISH) on metaphase chromosome spreads (Figure 1B). Whole-genome sequencing coupled with AmpliconArchitect analysis showed that these isogenic clones bore nearly identical *Kras* amplicon structures (Figure S1). Allele frequency analysis showed that the majority of *Kras* amplicons harbored the G12D mutation, suggesting a positive selection for the mutant *Kras* allele (Figure S2A). AmpliconArchitect additionally revealed that the *Myc* locus was also amplified on another ecDNA species, which was subsequently verified by *Myc* DNA FISH, showing *Myc* ecDNA or HSR amplification, respectively, in the EC1/2 or HSR1/2 clones (Figure 1B, Figure S1). The *Kras* and *Myc* amplification status of these lines was stably maintained in orthotopic tumors, as confirmed by DNA FISH on metaphase spreads (Figures 1C-D).

To investigate the transcriptional profile of these clones, we performed RNA-seq and found that EC1/2 and HSR1/2 exhibited similar phenotypes *in vitro*. Analyses of principal components and differentially expressed genes based on RNA sequencing data revealed similar transcriptional patterns among individual clones within the EC or HSR group, with only 1% of the transcriptome contributing to the difference between EC and HSR (Figures S2B-C). Gene set enrichment analysis (GSEA) demonstrated an insignificant difference in the KRAS signaling, and only one of the MYC signaling signatures was marginally upregulated in EC clones (Figure S2D). Moreover, all four clones showed negligible differences in cell proliferation capacity *in vitro* (Figure S2E). These data suggest that the variation of *Kras* and *Myc* copy number among clones at the bulk-cell level does not drive a significant phenotypic contrast *in vitro*.

However, tumors derived from EC1/2 cells grew more rapidly than those from HSR1/2 cells in immunocompetent C57BL/6J mice (Figure 1E). Consequently, mice bearing EC tumors exhibited significantly shorter survival times (Figure 1F). These results show that ecDNA-containing cancer cells promote more aggressive tumor progression than their isogenic HSR counterparts, likely through interactions with the TME rather than solely cell-intrinsic mechanisms.

### ecDNA-driven PDAC rapidly establishes an immunoevasive TME

Next, we investigated TME differences between EC and HSR tumors. We established orthotopic tumors using EC1 or HSR1 cells and harvested them at early- and mid-stages (10- and 15-days post-inoculation, respectively) (Figure 2A). To characterize the TME, we performed scRNAseq on mid-stage tumors from both groups. At this time point, tumor weights showed no significant difference (Figure S3A), thereby avoiding artifacts caused by size disparities or necrosis associated with larger tumor volumes. To minimize sampling bias, two bisected tumors within each group were randomly combined. A total of 3 samples for each group were dissociated to prepare single-cell suspensions (Figure 2A). Following enzymatic digestion, we captured approximately 20,000 cells per tumor type. Reference-based annotation using SingleR identified the cell composition of the TME,^24^ including cancer cells, immune cells, fibroblasts, and endothelial cells (Figure 2B), whose identities were verified using canonical biomarkers (Figure S3B).

**Figure 2.**
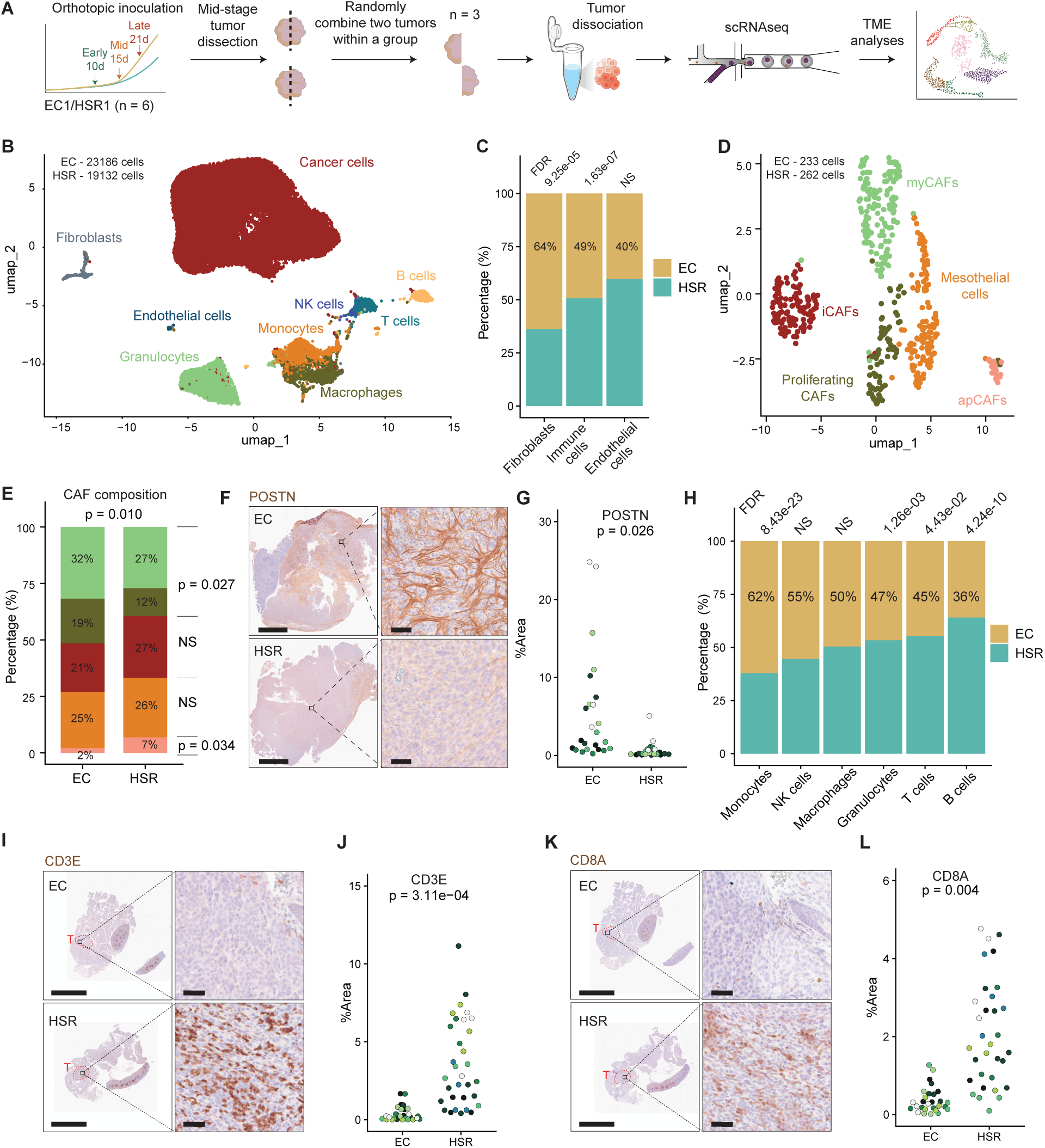
scRNAseq profiling of ecDNA and HSR-driven PDAC TME with IHC validation. (A) Schematic overview of the scRNAseq workflow. (B) UMAP visualization of all cells identified by scRNAseq from EC1 and HSR1 tumors. (C) Cell composition analysis of stromal and immune cell populations. Fisher’s exact test with FDR correction. NS indicates not significant. (D) UMAP visualization of CAF subclusters. (E) Cell composition analysis of CAF subclusters. Colors correspond to those in Figure 2D. Fisher’s exact test for the overall p-value, pairwise Fisher’s exact test with Benjamini-Hochberg correction for specific cell types. (F) Representative images of POSTN IHC staining showing both low- and high-magnification views. Scale bars 2.5 mm (low magnification), 50 μm (high magnification). (G) Quantification of POSTN IHC staining. Each color within a group represents an individual mouse. Four non-overlapping regions from the same section were analyzed and shown in the same color. Sample sizes: EC, n = 6; HSR, n = 6. Wilcoxon test. (H) Cell composition analysis of all *Ptprc*+ immune cells. Fisher’s exact test with FDR correction. NS indicates not significant. (I) Representative images of CD3E IHC staining showing low- and high-magnification views. Scale bars: 5 mm (low magnification) and 50 μm (high magnification). Red letter T and circles indicate the tumor area in the pancreas. (J) Quantification of CD3E IHC staining. Each color within a group represents an individual mouse. Four non-overlapping regions from the same section were analyzed and shown in the same color. Sample sizes: EC, n = 7; HSR, n = 8. Wilcoxon test. (K) Representative images of CD8A IHC staining showing low- and high-magnification views. Scale bars: 5 mm (low magnification) and 50 μm (high magnification). Red letter T and circles indicate the tumor area in the pancreas. (L) Quantification of CD8A IHC staining. Each color within a group represents an individual mouse. Four non-overlapping regions from the same section were analyzed and shown in the same color. Sample sizes: EC, n = 7; HSR, n = 8. Wilcoxon test.

Focusing on stromal and immune cell populations, EC tumors contained significantly more fibroblasts and slightly fewer immune cells compared to HSR tumors (Figure 2C). To characterize fibroblast heterogeneity, we extracted and re-clustered all fibroblasts from the dataset. Using established cancer-associated fibroblasts (CAFs) subtype markers, we identified inflammatory CAFs (iCAFs), myofibroblastic CAFs (myCAFs), antigen-presenting CAFs (apCAFs), proliferating CAFs (a proliferative myCAF subset), and mesothelial cells (Figures 2D, Figure S3C).^25–28^ EC tumors exhibited increased proportions of myCAFs and proliferating CAFs but decreased apCAFs (Figure 2E). As CAF networks are particularly resistant to enzymatic digestion due to dense extracellular matrix, our scRNAseq may underestimate CAF abundance. We therefore validated these findings using immunohistochemistry (IHC) for periostin (POSTN), a myCAF marker, confirming significantly higher myCAF composition in EC tumors (Figures 2F-G).

Within the immune cell populations, EC tumors contained fewer T and B cells (Figure 2H). We extracted and re-clustered all T cells, identifying CD8+ T cells and Tregs based on *Cd4*, *Foxp3*, and *Ctla4* expression (Figures S3D-E). ^29^ EC tumors exhibited reduced CD8+ T cells but increased Tregs, indicating a more immunosuppressive TME (Figure S3F). However, total T cell abundance was scarce in both EC and HSR tumors (<2.6% of TME cells), indicating that both groups exhibited an immune-cold phenotype by mid-stage (Figure S3G). To assess the onset of immunoevasion, we examined early-stage tumors (10 days post-inoculation) by IHC for CD3E and CD8A. HSR tumors showed abundant T cell infiltration, while EC tumors already exhibited significantly reduced T cell infiltration (Figures 2I-L). These findings indicate that EC tumors establish an immunoevasive TME rapidly after tumor initiation, preceding the immune-cold transition observed in HSR tumors.

### ecDNA heterogeneity drives *Kras* super-expressors

Next, we investigated the mechanism by which ecDNA-driven PDAC shapes an immunoevasive TME through myCAF expansion and T cell suppression. In contrast to focal gene amplification on chromosomes, ecDNAs drive oncogene copy number heterogeneity through unequal mitotic segregation, thereby generating a pool of genetically diverse cancer cells for selection.^30^ While cancer heterogeneity has been linked to aggressive phenotypes, the mechanisms by which ecDNAs foster a more immunoevasive TME remain unclear. We hypothesized that a small subset of ecDNA-high or ecDNA-low cell populations may have a distinct transcriptional profile, which altered communication between cancer cells and host cells.

Therefore, we profiled the *Kras* expression level for EC and HSR tumors. Among cancer cells, the lower bound of the *Kras* expression was comparable between EC and HSR tumors; however, EC cancer cells exhibited a markedly higher upper bound and greater variation of *Kras* expression (Figure 3A), as assessed by the median absolute deviation (MAD) and interquartile range (IQR). This pattern was consistently observed at the levels of DNA copy number and protein abundance, with *Kras* DNA FISH and protein immunofluorescence (IF) signals showing elevated upper bounds and greater variation in EC tumors (Figures S4A-D). Co-staining of *Kras* DNA FISH and RAS protein IF revealed a positive correlation between *Kras* ecDNA copy number and protein abundance (Figures S4E-F), indicating that expression variation of *Kras* was dictated by ecDNA copy number heterogeneity.

**Figure 3.**
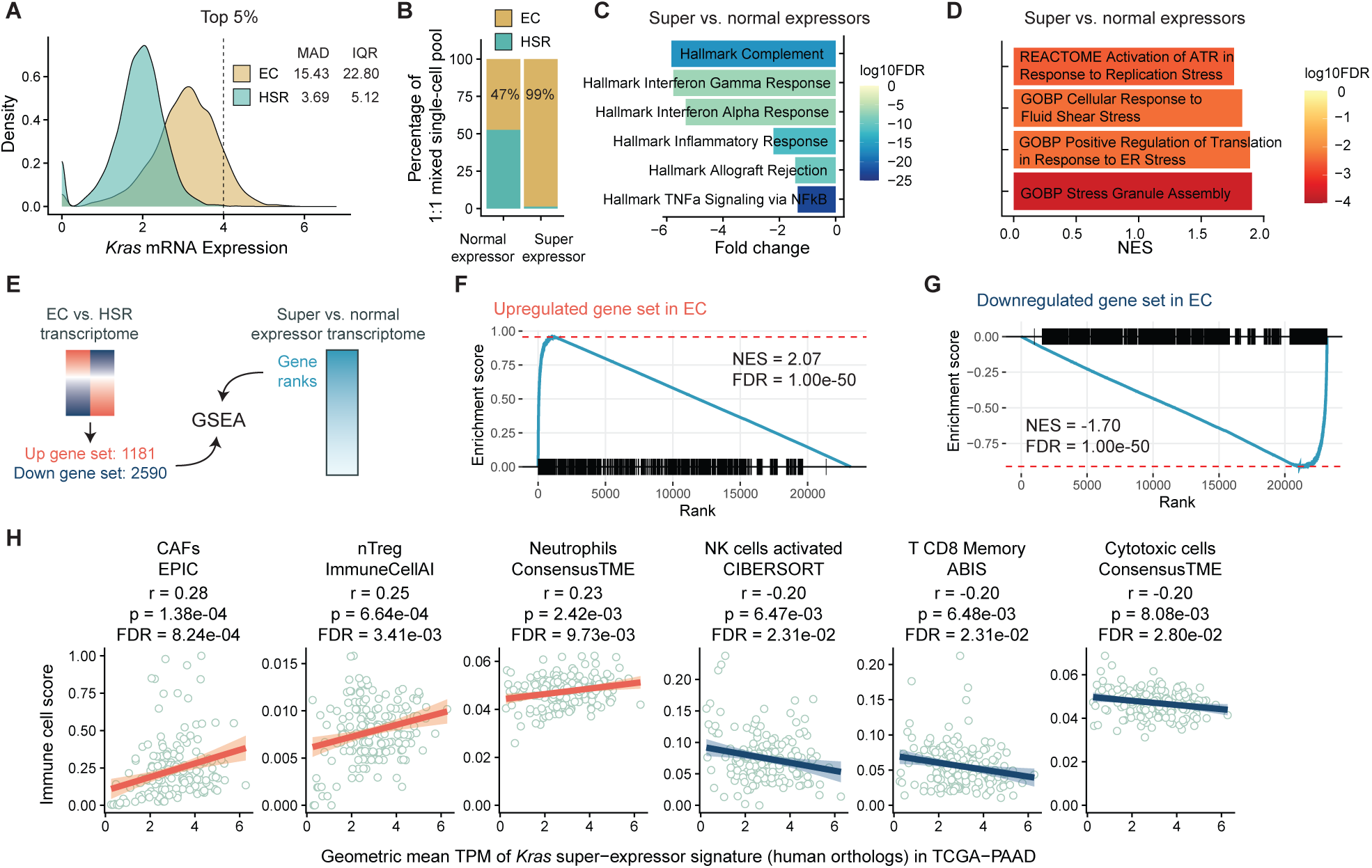
Molecular characteristics of ecDNA-driven *Kras* super-expressors. (A) Density plot of *Kras* mRNA expression of EC and HSR cancer cells from scRNAseq data. The distribution variance was assessed using median absolute deviation (MAD) and interquartile range (IQR). The vertical dashed lines indicate the 95% percentile, which was used to distinguish super- and normal-expressors. Number of cells analyzed: EC, n = 10,000; HSR, n = 10,000. (B) Proportion analysis of normal-expressor and super-expressor in EC and HSR groups based on the same subset of cells in Figure 3A. (C) Differential pathway analysis using SCPA with the mouse-ortholog hallmark gene set as input. (D) Differential pathway analysis using GSEA with the stress-related pathways from mouse collections as input. (E) Schematic showing the input of GSEA analysis. The DEGs between EC and HSR transcriptomes were used to generate two gene sets: the Upregulated gene set in EC and the Downregulated gene set in EC. A ranked list of genes was generated based on expression differences between super- and normal-expressors. (F-G) GSEA results show the contrast between *Kras* super- and normal-expressors resembles the differences between EC and HSR cancer cells. (H) Correlation between the expression of the human orthologs of *Kras* super-expressor signature genes and the immune cell infiltration scores in the TCGA pancreatic cancer (TCGA-PAAD) cohort from TIMEDB, analyzed using the Pearson correlation test with FDR correction.

In contrast, the variance of *Myc* expression did not show a substantial difference between EC and HSR cancer cells (Figure S4G). Furthermore, DNA FISH-IF analysis revealed a weaker correlation between *Myc* ecDNA copy number and protein abundance (Figures S4H-I). Notably, cancer cells with the highest MYC protein expression did not possess the highest *Myc* ecDNA copy number, suggesting that additional regulatory mechanisms beyond DNA copy number contribute to *Myc* expression in this cancer model.

Taken together, these data suggest that *Kras* ecDNA may give rise to a subpopulation of cancer cells with exceptionally high *Kras* expression, hereinafter referred to as *Kras* super-expressors.

### *Kras* super-expressors are associated with an immunoevasive TME

To investigate the functional role of ecDNA-driven *Kras* super-expressors, we performed pathway analysis using scRNAseq data (Figure S5A). An equal number of cancer cells (n = 10,000) was randomly sampled and pooled from EC and HSR tumors. Super- and normal-expressors were defined based on above or below the top 5% *Kras* expression level cutoff (Figure 3A, Figure S5B). Under this threshold, KRAS signaling was significantly upregulated in super-expressors compared to normal-expressors within EC tumors, whereas normal-expressors between EC and HSR tumors showed no significant difference in KRAS signaling (Figure S5B). This strategy optimized discovery sensitivity while minimizing false positives.

Under this definition, the vast majority (99%) of *Kras* super-expressors in the 1:1 mixed single-cell pool originated from EC tumors (Figure 3B, Figure S5C). Single-cell pathway analysis using Single cell pathway analysis (SCPA) revealed that *Kras* super-expressors downregulated numerous immune response pathways,^31^ including the interferon alpha and gamma pathways, as well as the TNF-alpha signaling (Figure 3C). However, these *Kras* super-expressors also experienced cellular stress, including replication stress and ER stress, as shown by GSEA (Figure 3D). For example, genes involved in single-stranded DNA binding proteins (*Rpa1*/*2*), the 9-1-1 complex (*Rad9a*/*Rad1*/Hus1), and the check1-claspin axis (*Chek1*/*Clspn*) were upregulated, indicating elevated replication stress (Figure S5D). Further analysis using *Kras* and *Myc* DNA FISH in EC tumors revealed that high copy numbers of *Kras* and *Myc* rarely coexisted within the same cancer cell (Figure S5E), suggesting a potential mutual exclusivity in ecDNA-driven amplification shaped by oncogene overdose.^32^ These findings suggest that, while *Kras* super-expressors may evade immune surveillance better, elevated cellular stress may limit their proliferation (Figure S5F).

To assess whether *Kras* super-expressors contribute to the phenotypic divergence between EC and HSR tumors, we performed GSEA to determine whether gene sets derived from differentially expressed genes between EC and HSR significantly changed between super- and normal-expressors transcriptomes (Figure 3E). The results revealed significant and concordant differences, supporting the hypothesis that ecDNA-driven *Kras* super-expressors shape the biological distinction between EC and HSR tumors (Figures 3F-G).

To further explore the impact of *Kras* super-expressors on the TME, we analyzed the correlation between the expression of *Kras* super-expressor signature genes and the immune cell score in the TCGA pancreatic cancer (TCGA-PAAD) cohort from TIMEDB.^33^ The top 30 signature genes (Figure S5A, Supplementary Table 1) were mapped to their human orthologs to obtain transcripts per million (TPM) values, and the geometric mean was calculated per subject. Correlation analysis revealed that the *Kras* super-expressor signature was associated with increased infiltration of CAFs, natural Tregs, and neutrophils, alongside decreased infiltration of CD8+ T cells and activated NK cells, mirroring the immunoevasive phenotype in EC tumors (Figure 3H). These data suggest that *Kras* super-expressors may play a pivotal role in establishing an immunoevasive TME.

### *Kras* super-expressors induce myCAF expansion via AREG signaling

To identify effector genes in *Kras* super-expressors that suppressed the immune response, we applied a multi-step filtering strategy. Candidate genes were selected based on three criteria: (1) upregulated in *Kras* super-expressors compared to normal-expressors; (2) upregulated in TCGA-PAAD tumors relative to normal pancreatic tissues from the Genotype-Tissue Expression (GTEx) dataset; (3) annotated as cell-cell communication ligands from CellChat (Figure 4A, Figure S5A).^34^ This approach yielded five candidate genes, including *AREG*, *CEACAM1*, *CSF2*, *PDCD1LG2*, and *PTHLH* (Figures 4A-B, Figures S6A-B). Among these, upregulation of *AREG* and *CSF2* was significantly associated with a shorter overall survival in the TCGA-PAAD cohort (Figure 4C, Figure S6B).

**Figure 4.**
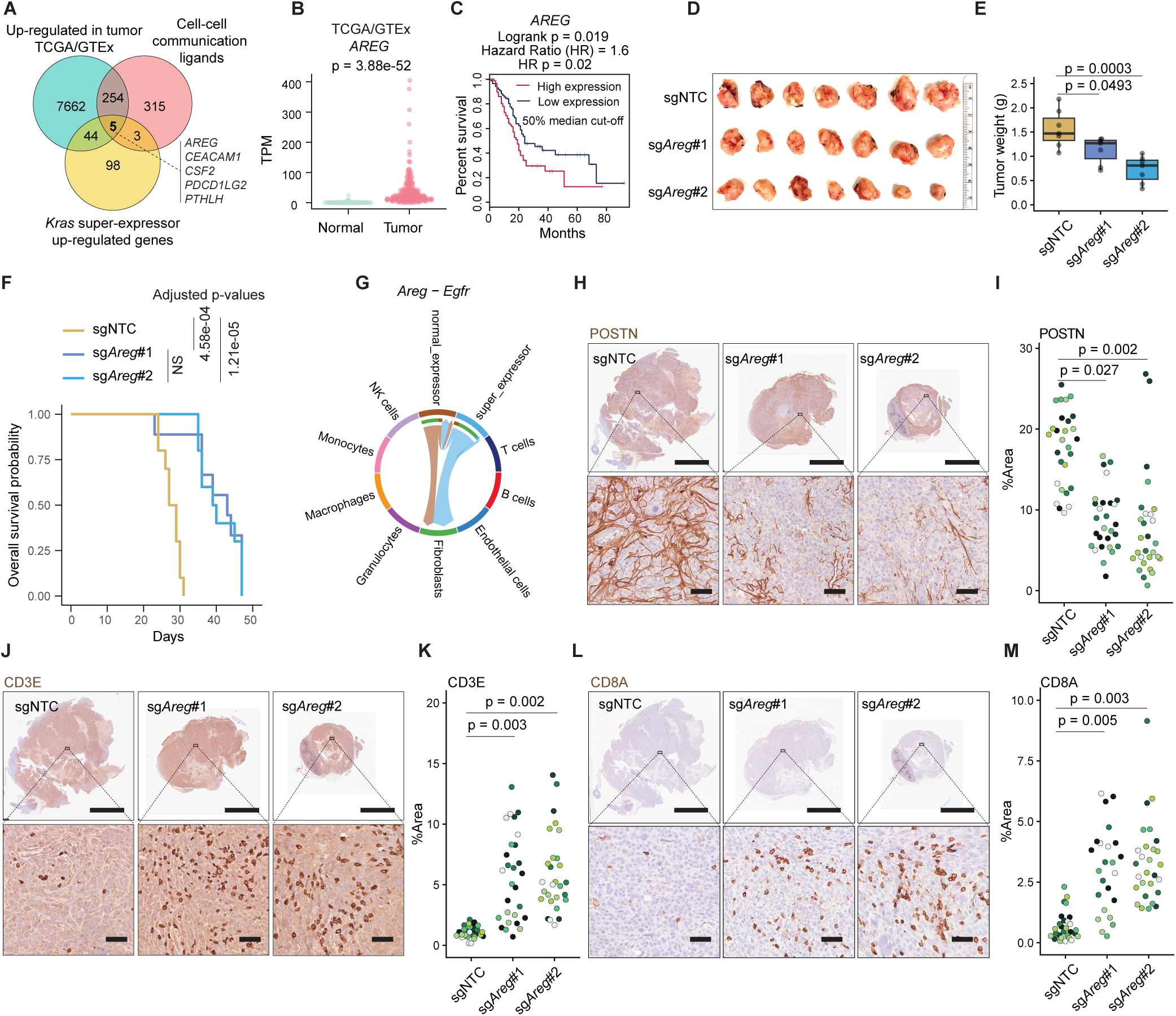
Functional validation of *Areg* in PDAC TME remodeling. (A) Venn diagram showing the identification of effector genes in *Kras* super-expressors. Upregulated genes between TCGA-PAAD tumors and GTEx normal pancreas samples were intersected with ligand genes from CellChat and *Kras* super-expressor upregulated genes from scRNAseq analysis. (B) Dot plot showing the differential expression of *AREG* between TCGA-PAAD tumor samples and GTEx normal samples. Wilcoxon test. (C) Overall survival of TCGA-PAAD cohort stratified by *AREG* expression (median 50% cutoff). Log-rank test. (D) Gross image of orthotopic tumors from C57BL/6J mice inoculated with CT1BA5EC1 cell lines with or without *Areg* knockout. NTC indicates non-targeting control. (E) Box plot showing the tumor weight difference among the three groups. Sample sizes: sgNTC, n = 7; sg*Areg*#1, n = 7; sg*Areg*#2, n = 7. One-way ANOVA with Tukey’s HSD. (F) Overall survival of C57BL/6J mice bearing CT1BA5EC1 tumors with or without loss of *Areg*. Sample sizes: sgNTC, n = 10; sg*Areg*#1, n = 9; sg*Areg*#2, n = 10. Pairwise log-rank test with FDR correction. NS indicates not significant. (G) Chord diagram generated by CellChat based on scRNAseq data showing the Areg-Egfr ligand-receptor communication among different cell types in EC samples. Each chord represents communication between two cell types. The direction of the chords is from the sender to the receiver. The width of a chord represents interaction strength. (H-M) Representative images of POSTN (H), CD3E (J), and CD8A (L) IHC staining with both low- and high-magnification views. Scale bars 5 mm (low magnification), 50 μm (high magnification). Dot plots show the quantification of POSTN (I), CD3E (K), and CD8A (M). Each color within a group represents an individual mouse. Four non-overlapping regions from the same section were analyzed and shown in the same color. Sample sizes: sgNTC, n = 7; sg*Areg*#1, n = 7; sg*Areg*#2, n = 7. Kruskal–Wallis test with Bonferroni correction.

*Csf2* (granulocyte-macrophage colony-stimulating factor, GM-CSF) is known to orchestrate an immunosuppressive PDAC TME by regulating myeloid-derived suppressor cells.^6,7^ *Areg* encodes amphiregulin, an EGF-like growth factor that has been implicated in promoting metastasis via autocrine from myCAFs in PDAC.^35^ We decided to focus on *Areg* because of its higher expression in our mouse models (Figures S6C-E).

To confirm that KRAS signaling regulates *Areg* expression, we performed RNAi-mediated *Kras* knockdown in EC1 cells, which significantly reduced *Areg* expression (Figure S6F). Consistently, pharmacological inhibition of the MEK-ERK pathway using trametinib (MEKi) and SCH772984 (ERKi) dramatically suppressed *Areg* expression (Figures S6G-J). Inhibition of the mTOR signaling by rapamycin (mTORC1i) and torin 2 (mTORC1/2i) yielded a significant yet milder suppression effect on *Areg* expression (Figures S6G-J).

To investigate the functional role of *Areg* in PDAC, we knocked out *Areg* in EC1 cells (Figure S7A). *Areg* deletion did not alter *Kras* or *Myc* ecDNA copy numbers, nor did it impair cell viability *in vitro* (Figures S7B-E). However, orthotopic implantation of *Areg*-deficient EC1 cells resulted in significantly reduced tumor growth and prolonged survival in mice (Figures 4D-F), suggesting that amphiregulin promotes PDAC malignancy through interactions with the TME rather than a cancer cell-intrinsic mechanism.

scRNAseq analysis showed that *Kras* super-expressors exhibited the highest transcription of *Areg*, whereas the amphiregulin receptor *Egfr* was primarily expressed in fibroblasts (Figure S7F). Cell-cell communication analysis further revealed a stronger ligand-receptor interaction between *Kras* super-expressors (sender) and fibroblasts (receiver) in EC tumors, even though super-expressors accounted for only 5% of the cancer cells (Figure 4G). These findings prompted us to investigate the role of *Areg* in myCAF expansion.

Knockout of *Areg* in EC1 cancer cells significantly reduced the number of myCAFs in EC tumors (Figures 4H-I). Moreover, the number of Ki67-positive, actively proliferative myCAFs also decreased in tumors derived from *Areg*-deficient EC1 cells (Figures S7G-H). These data suggest that tumor-cell-derived amphiregulin is a crucial factor in promoting myCAF proliferation in EC tumors.

Given the established role of myCAFs in suppressing T cell function in PDAC,^36,37^ we examined the relationship between myCAF abundance and immune infiltration in human tumors. In the TCGA-PAAD cohort, the myCAF signature score, defined as the geometric mean expression of established myCAF marker genes (Supplementary Table 2), was negatively correlated with the infiltration of multiple cytotoxic immune cell populations, including CD4+ naive T cells, CD8+ T cells, Th1 cells, and natural killer T (NKT) cells (Figure S7I). To test whether cancer cell-derived amphiregulin contributes to immune evasion, we examined T cell infiltration in tumors derived from *Areg*-deficient EC1 cells. Consistent with reduced myCAF abundance, genetic ablation of *Areg* significantly increased CD3+ and CD8+ T cell infiltration (Figures 4J-M). Notably, this effect persisted even in late-stage tumors (21 days post-orthotopic injection, as shown in Figures 4J-M), when both EC and HSR tumors typically exhibit profound immune evasion.

Collectively, these data demonstrate that *Kras* super-expressors shape an immunoevasive TME by delivering amphiregulin, which promotes myCAF expansion and suppresses T cell infiltration.

### Clonal expansion of *KRAS*-high cells fosters an immunoevasive spatial niche

During PDAC progression in *KPfC* mice, we observed sporadic clusters of cancer cells exhibiting high *Kras* DNA FISH signals with morphological features resembling ecDNA (Figure S8). This finding suggests a multi-clonal emergence and expansion of *Kras* ecDNA-driven cancer cells, leading us to investigate whether clonal expansion of *KRAS*-high cells contributes to the formation of a spatially organized immunoevasive niche in human samples.

To dissect the spatial organization of *KRAS*-high and *KRAS*-low cancer cell populations and their associated tumor microenvironments at single-cell resolution, we employed the Xenium *in situ* platform for high-plex spatial transcriptomics. We designed a custom 480-gene panel composed of established markers of cancer cells, immune cells, and CAF subtypes (Supplementary Table 3). We used it to profile six human PDAC specimens. In one specimen, a distinctive, heterogeneous *KRAS* expression pattern in cancer cells was identified (Figure S9A). Unsupervised clustering of this specimen identified the following major cell types: cancer cells, myCAFs, apCAFs, myeloid cells, T cells, endothelial cells, and mast cells (Figures 5A-B, Figure S9B).

**Figure 5.**
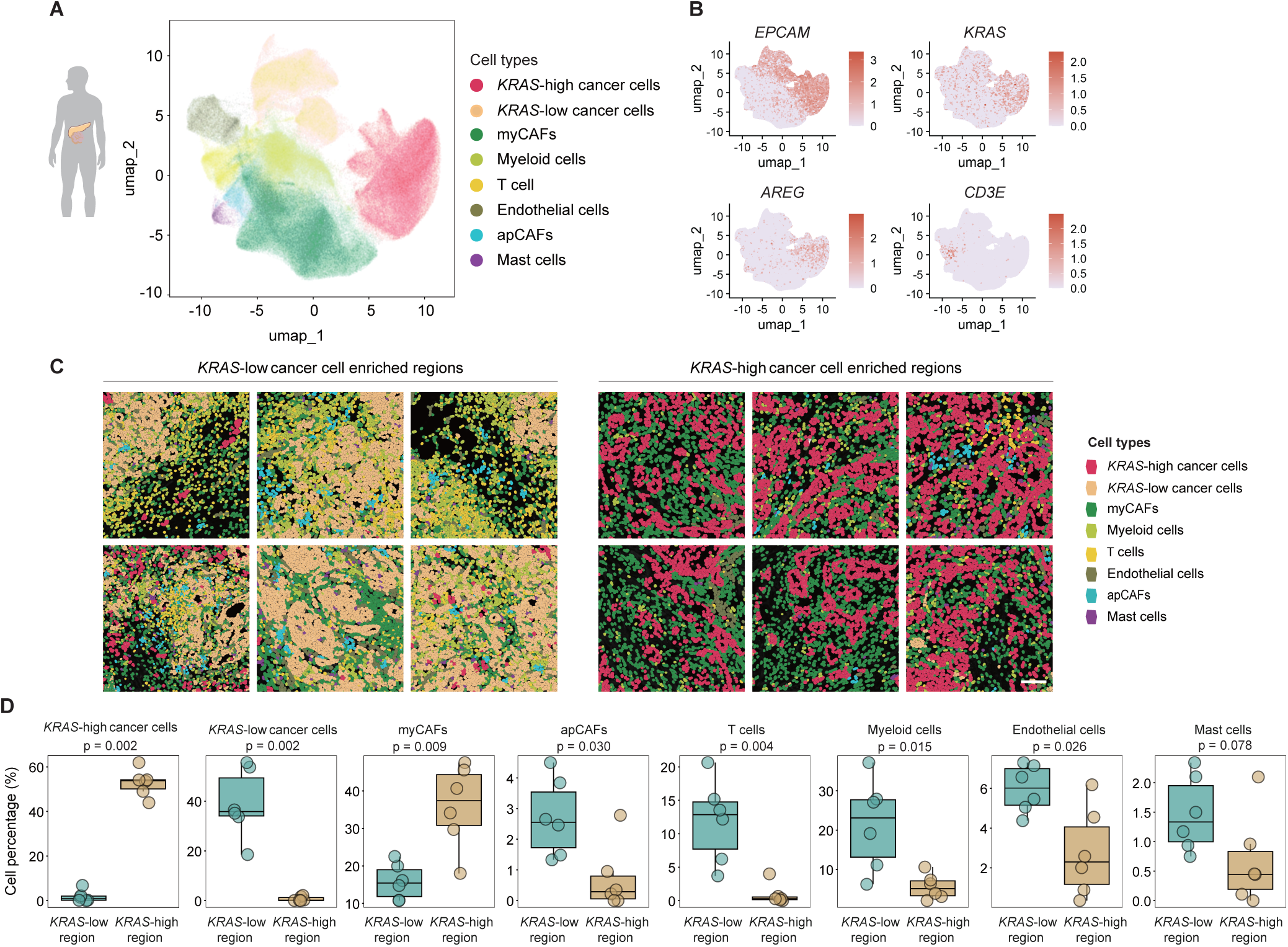
Spatial analysis of *KRAS*-high and *KRAS*-low niches in PDAC. (A) UMAP visualization of major cell populations identified by Xenium spatial transcriptomics in a human PDAC sample. (B) UMAP visualization of *EPCAM*, *KRAS*, *AREG,* and *CD3E* expression in the human PDAC tissue of the Xenium spatial transcriptomic data. (C) Spatial distribution of major cell populations in representative tumor regions enriched for *KRAS*-high versus *KRAS*-low cancer cells. Cell types are color-coded as indicated. Number of regions: *KRAS*-high, n=6; *KRAS*-low, n=6. Scale bar: 50 μm. (D) Box plots displaying the proportional distribution of cell populations in spatial regions of *KRAS*-high and *KRAS*-low cancer cells. Number of regions: *KRAS*-high, n=6; *KRAS*-low, n=6. Wilcoxon test.

We found that cancer cells were divided into two distinct clusters, characterized by differential *KRAS* expression levels (Figure 5A, Figure S9B). Notably, *KRAS*-high cancer cells also exhibited elevated levels of *AREG*, recapitulating the results observed in mouse tumors (Figure 5B, Figure S9B). Spatial visualization confirmed that *AREG*-expressing cells were predominantly localized within *KRAS*-high cancer cell regions (Figure S9C).

To characterize the stromal composition of these distinct tumor subpopulations, we quantified CAF and immune cell densities in spatially defined regions enriched for either *KRAS*-high or *KRAS*-low cancer cells (Figure S9A). Consistently, *KRAS*-high regions exhibited significantly higher myCAF abundance and reduced T cell infiltration compared to *KRAS*-low regions (Figures 5C-D, Figure S9C). These data demonstrate that the KRAS-AREG-axis mediated immunoevasive program, defined by myCAF enrichment and T cell exclusion, manifests in human PDAC as spatially distinct tumor microenvironments, driven by cancer cell-intrinsic heterogeneous KRAS expression.

## Discussion

Alterations in oncogenes and tumor suppressor genes are central to TME remodeling^1^. Although ecDNA amplification has been associated with immune evasion^12–15,18^, its causal role remains unclear. Here, we show that ecDNA-driven amplification of *Kras* generates a small subpopulation of cancer cells with extremely high copy numbers. These *Kras* super-expressors downregulate immune response pathways and secrete amphiregulin (AREG), which establishes an immunoevasive niche characterized by enriched myCAF and reduced T cell infiltration. Although we do not exclude the possibility that a pre-existing cold TME favors ecDNA formation, our data provide direct evidence that ecDNA-mediated heterogeneity actively shapes the TME.

Acentric ecDNA drives intratumoral heterogeneity through non-Mendelian segregation, producing daughter cells with variable ecDNA copy numbers.^9,38^ This contributes to therapy resistance and poor prognosis.^10,11,39–41^ Our study advances current understanding by demonstrating that, in the *KPfC* mouse PDAC model, ecDNA-driven *Kras* super-expressors act as powerful architects of an immunoevasive TME. In human PDAC, heterogeneous *KRAS* expression similarly correlates with spatial variation in myCAF and immune cell organization, where cancer cells with elevated KRAS levels are spatially associated with a more immunoevasive niche. However, excessive oncogene signaling, such as hyperactivation of the KRAS-ERK pathway, can impair cell fitness via senescence and apoptosis.^42^ Signature analysis of our scRNAseq data reveals elevated cellular stress, including replication stress, in *Kras* super-expressors, explaining why ecDNA copy number cannot increase indefinitely. Notably, ecDNA enables rapid adaptation through asymmetric segregation, allowing daughter cells to resolve oncogene overdose more easily than chromosomal counterparts.

Comparative analysis of ecDNA and HSR-driven expression reveals greater variation in *Kras* ecDNAs, but not in *Myc* ecDNAs. The weaker correlation between *Myc* ecDNA copy number and gene expression suggests additional regulatory mechanisms modulate *Myc* transcription, indicating that ecDNA copy number heterogeneity does not always translate into transcriptional heterogeneity. Furthermore, PDAC cells rarely harbor extremely high copies of both *Kras* and *Myc* ecDNAs, suggesting the presence of post-segregation selection pressures that refine ecDNA composition beyond co-segregation.^43^

Myofibroblastic CAFs represent a critical barrier to anti-tumor immunity in PDAC.^36,44,45^ These activated fibroblasts not only generate dense extracellular matrices that physically exclude immune cells from the tumor parenchyma, but also actively secrete immunosuppressive factors that impair T cell function. The abundance of myCAFs has been reported to be inversely associated with anti-tumor immunity,^45^ underscoring the importance of understanding the molecular mechanisms that drive myCAF expansion in PDAC. Our study reveals that ecDNA-driven *KRAS* super-expressors actively remodel the TME by promoting myCAF proliferation, thereby establishing an immunoevasive niche that facilitates tumor progression.

Integrating scRNAseq, spatial transcriptomics, and functional genetic data, we identify *AREG* as a critical mediator of *KRAS* super-expressor-driven myCAF expansion and subsequent immune exclusion. Previous studies have demonstrated that autocrine AREG signaling in myCAFs promotes metastasis by activating EGFR-mediated motility and invasiveness.^35^ Our findings reveal a complementary paracrine mechanism in which cancer cells, specifically those with extremely high *KRAS* expression driven by ecDNA heterogeneity, secrete elevated levels of AREG to promote myCAF proliferation in the surrounding stroma, thereby excluding T cells. These data position AREG as a critical regulator during PDAC progression.

While *AREG* emerges as a central player, other effector genes, including *CEACAM1*, *CSF2* (GM-CSF), *PDCD1LG2* (PD-L2), and *PTHLH*, have also been implicated in PDAC pathology.^6,46–48^ A comprehensive mapping of the ecDNA-mediated crosstalk landscape is required to fully understand the interaction of ecDNA-driven cancer cells and TME. In addition, the mechanisms underlying *Kras* and *Myc* ecDNA formation in PDAC remain unclear. The clonal organization of *Kras*-amplified cancer cells suggests that ecDNA genesis may require additional genetic events beyond *Trp53* deficiency.^13^ Future study is warranted to identify these prerequisites for ecDNA pathogenesis and their roles in tumor evolution.

In summary, our findings highlight ecDNA as a dynamic genetic element that not only amplifies oncogene expression but also actively remodels the tumor ecosystem to promote immune evasion. This underscores the importance of incorporating ecDNA biology into therapeutic strategies targeting both cancer cell-intrinsic and microenvironmental vulnerabilities.

## STAR Methods

### Key resources table

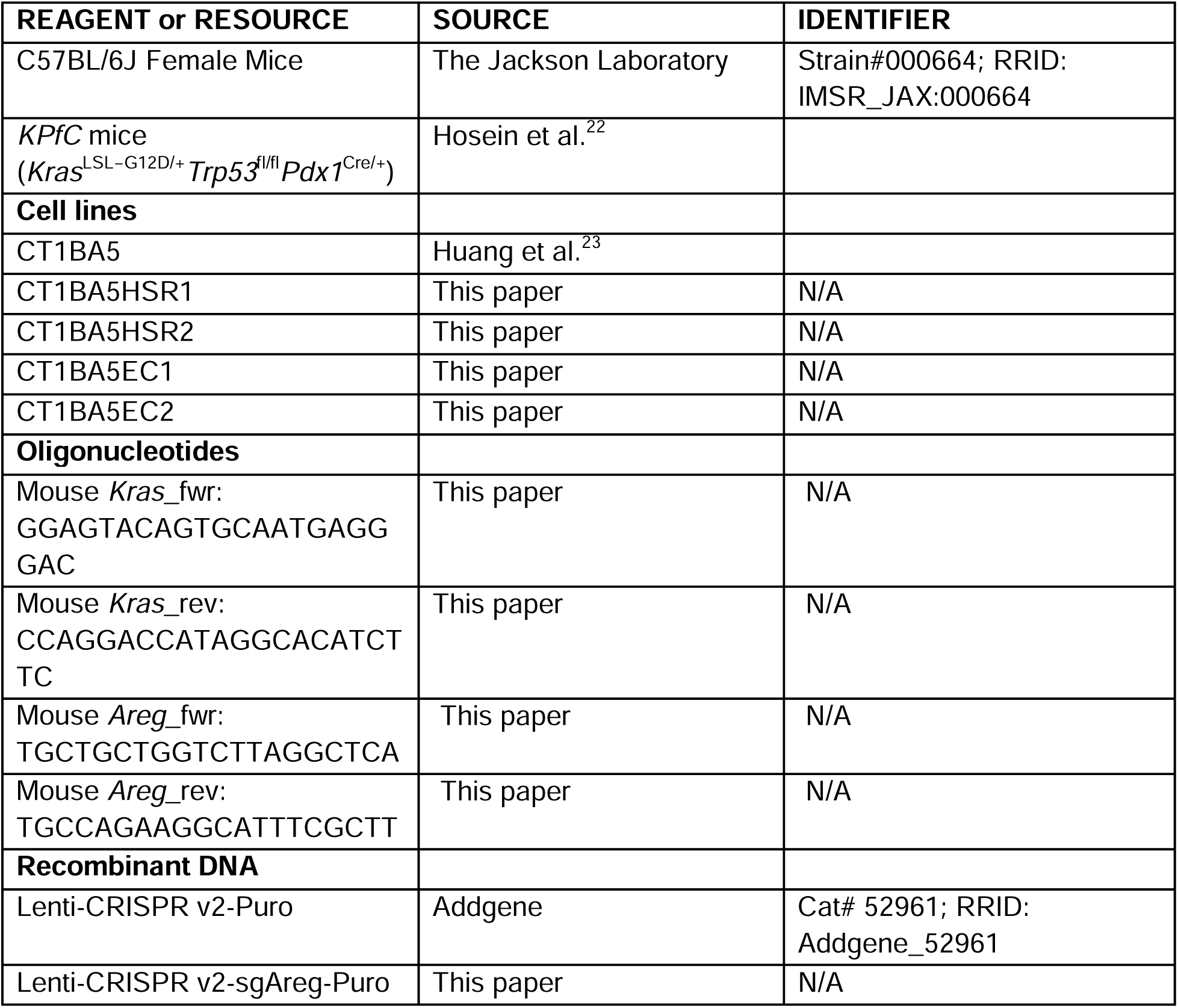
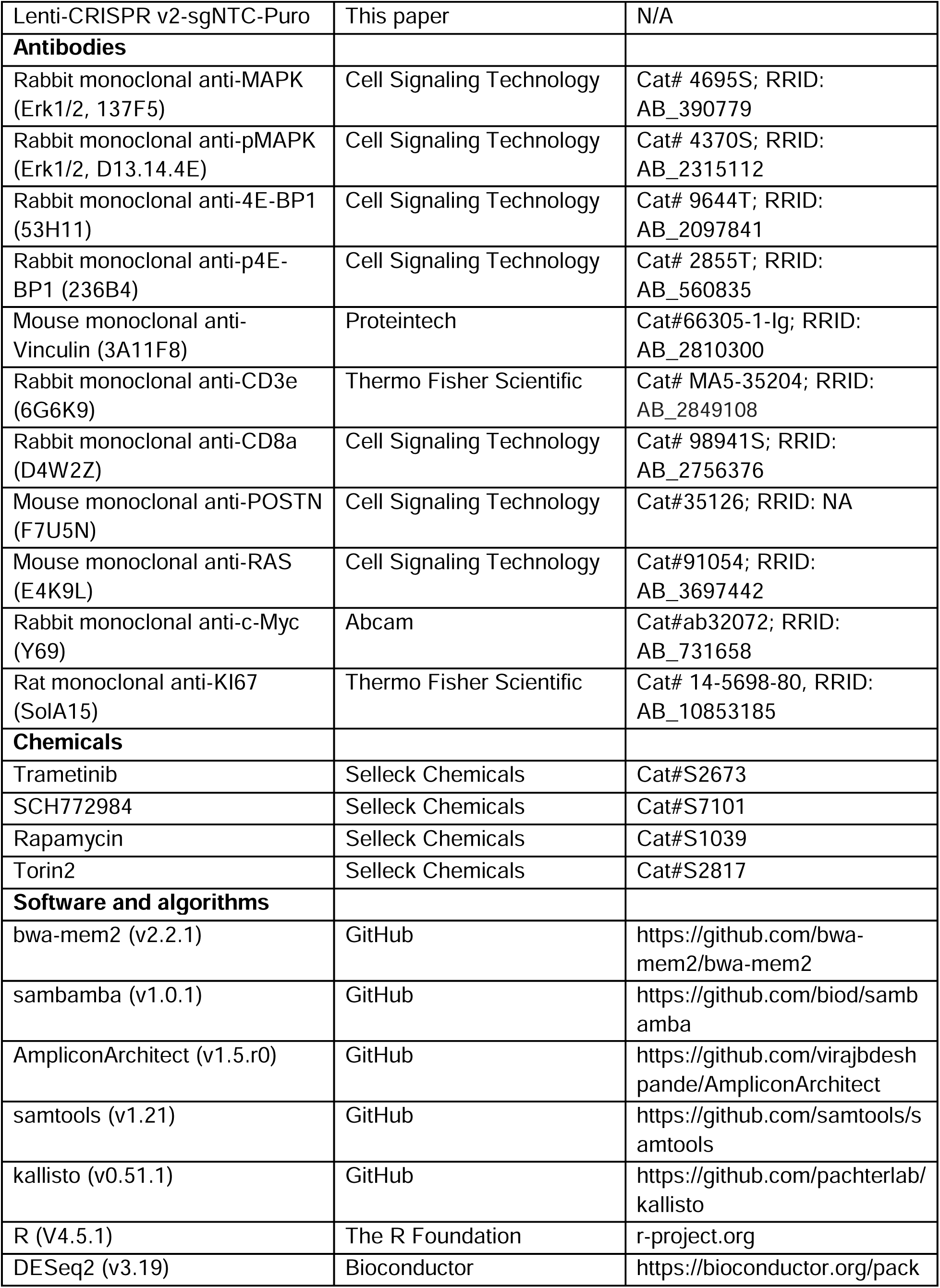
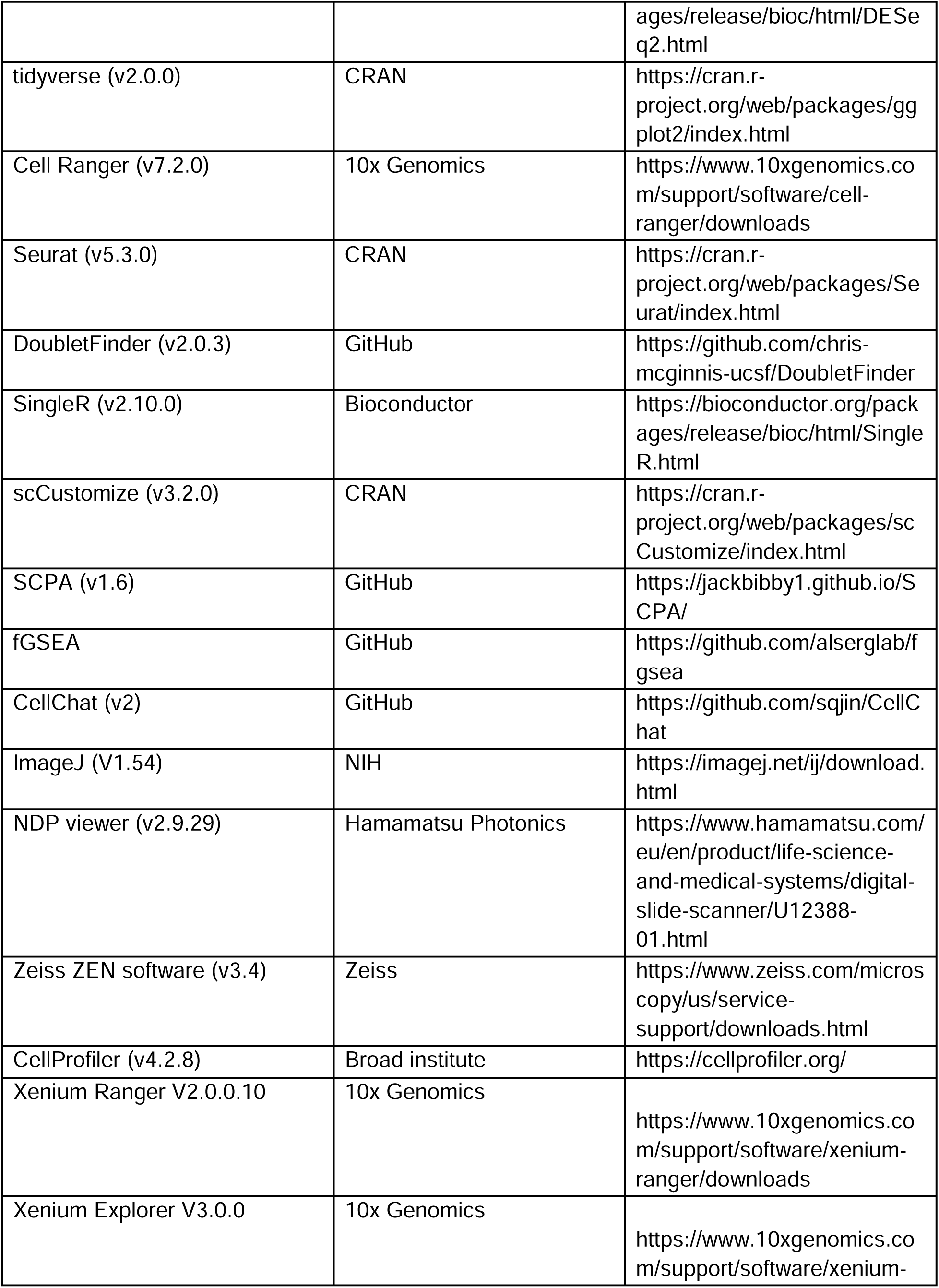
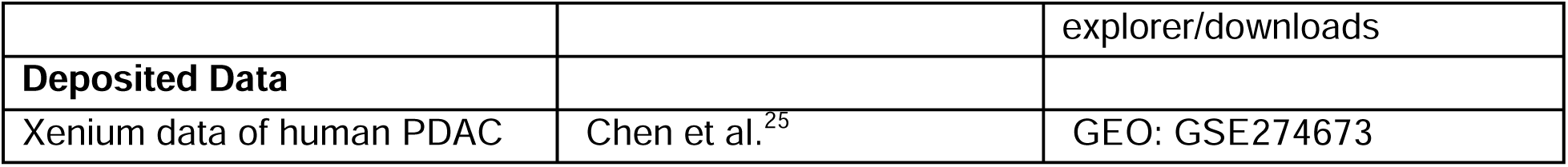

### Experimental model and study participant details

#### Mice

*KPfC* (*Kras^LSL-G12D/+^*; *Trp53^fl/fl^*; *Pdx1^Cre/+^*) mice were generated as previously described.^22^ The KPfC mouse had a pure C57BL/6 genetic background. All other mice used in this study were purchased from The Jackson Laboratory (6–9-week-old female C57BL/6J, strain #000664). All mice were maintained under specific pathogen-free conditions in the animal facility at the University of Texas Southwestern Medical Center at Dallas, in accordance with protocols approved by the Institutional Animal Care and Use Committee (IACUC).

#### Cell lines

CT1BA5 cells were established as previously described from a female *KPfC* mouse PDAC model (*Kras^LSL-G12D/+^*; *Trp53^fl/fl^*; *Pdx1^Cre/+^*) on a C57BL/6 background.^23^ Cells were cultured in Dulbecco’s Modified Eagle’s Medium/F12 (DMEM/F12; Corning, 10-092-CV) supplemented with 10% Fetal Bovine Serum (FBS; Corning, 35-011-CV) and 1× Penicillin-Streptomycin (Corning, 30-002-CI) at 37°C with 5% CO_2_. All cells were regularly tested for mycoplasma.

#### Patient samples

Human PDAC specimens were procured from the UT Southwestern Tissue Management Shared Resource following informed patient consent and approval by the UT Southwestern Institutional Review Board. Formalin-fixed paraffin-embedded (FFPE) tissue blocks were sectioned at 5-μm thickness. Histopathological diagnosis was verified by an experienced pathologist through hematoxylin and eosin staining of representative sections. Adjacent serial sections from these blocks were subsequently utilized for spatial transcriptomic profiling. Patient demographic information, including age and sex, was not disclosed to maintain confidentiality.

### Method details

#### Tissue dissociation

Tumor samples were minced into 1-2 mm^3^ pieces on ice and transferred to 15 mL tubes containing warmed 10 mL tumor digestion buffer (9 mL DMEM + 1 mL 10× digestive buffer; see recipe below), then incubated at 37°C with shaking at 100 rpm for 30 minutes. The resulting suspensions were filtered through a 70-µm cell strainer, and enzymatic activity was quenched by adding ice-cold medium containing 10% fetal bovine serum (FBS). The suspensions were spun down and washed once using ice-cold medium with 10% FBS. To lyse blood cells, the cell pellet was resuspended gently by adding 5 mL ice-cold ACK lysing buffer (Thermo Fisher Scientific, A1049201) and incubated for 10 min on ice. Then ice-cold DPBS (Corning, 21031CV) with 0.04% BSA was added to the cells, and the cells were spun down at 300 ×g for 5 minutes at 4°C.

The 10× digestive stock buffer contains collagenase type I (450 units/ml, Worthington, LS004214), collagenase type II (150 units/mL, Worthington, LS004202), collagenase type III (450 units/mL, Worthington, LS004206), collagenase type IV (450 units/mL, Worthington, LS004210), elastase (0.8 units/mL, Worthington, LS002290), hyaluronidase (300 units/mL, Sigma, H3506100MG), and DNase type I (250 units/mL, Sigma-Aldrich, D452710KU) in PBS.

#### Metaphase chromosome spread

Cells were treated with KaryoMAX (Gibco, 15212012) for 3 hours, and the single-cell suspension was harvested by trypsinization and centrifugation at 500 ×g for 5 minutes. After washing with PBS, cells were resuspended in 600 µL of 75 mM KCl and incubated at 37°C for 25 minutes, followed by fixation with 600 µL freshly prepared Carnoy’s fixative solution (3:1 Methanol: Glacial acetic acid). Cells were spun down at 800 ×g for 2 minutes. After another three fixations, cells were resuspended in 300 µl fixative solution, dropped onto the humidified slide, and air-dried for DAPI staining at room temperature. Finally, the slides were mounted with antifade mounting medium (Vector Laboratories, H-1000-10) and sealed with nail polish.

#### OligoPaint DNA FISH on metaphase chromosome spreads

Air-dried slides with metaphase chromosome spreads were equilibrated in 2× SSC and dehydrated in 70%, 85%, and 100% ethanol for 2 minutes each. Primary OligoPaint *Kras* and *Myc* probes diluted in hybridization buffer (1:4) were applied to slides and mounted with coverslips. Slides were denatured for 2 minutes at 75°C and hybridized overnight at 37°C in a humidified chamber. Next, the slides were washed twice with 0.4× SSC containing 0.3% IGEPAL for 10 minutes at 45°C, followed by a single wash with 2× SSC containing 0.1% IGEPAL for 10 minutes at room temperature. Slides were then carefully dried before applying the secondary OligoPaint probe labeled with a fluorophore, diluted in hybridization buffer (1:4), and incubating at 37°C for 1 hour in the dark. Post-hybridization washes were repeated as above. Finally, the slides were stained with DAPI and mounted using mounting medium (Prolong Diamond Antifade Mountant, Thermo Fisher Scientific, P36961).

OligoPaint pool was designed and synthesized as previously reported.^49^ In brief, approximately 1,000 oligonucleotides, each 68 to 80 bases long, were designed to span a 200-kb genomic region using OligoMiner software. Diverse two 20-nt adapters were added to the 5’ and 3’ ends to enable PCR, *in vitro* transcription, reverse transcription, and fluorescent secondary probe binding. The following genomic coordinates were used for OligoPaint pool design: Mouse *Myc*, chr15:61,950,801-62,150,800; Mouse *Kras*, chr6: 45,129,701-145,329,700.

#### Whole genome sequencing (WGS)

WGS libraries were prepared using NEBNext Ultra II DNA Library Prep Kit for Illumina (NEB, E7645L) following the manufacturer’s manual. Sequencing was performed in paired-end 150 bp (PE150) mode. Raw sequencing data were aligned to the mm10 mouse reference genome using bwa-mem2 (v2.2.1).^50^ Aligned reads were sorted and deduplicated with sambamba (v1.0.1),^51^ then analyzed with AmpliconArchitect (v1.5.r0) to extract ecDNA structure and copy number.^52^ The *Kras* G12D mutation frequency was analyzed with the mpileup function in samtools (v1.21).^53^

#### Bulk-cell RNA sequencing (RNAseq)

RNAseq libraries were prepared using the NEBNext Ultra II RNA Library Prep Kit for Illumina (NEB, E7770L) with the NEBNext Poly(A) mRNA Magnetic Isolation Module (NEB, E7490) following the manufacturer’s manual. Sequencing was performed in paired-end 150 bp (PE150) mode. Raw sequencing data were analyzed using kallisto (v0.51.1) with the mm39 mouse reference genome.^54^ Differentially expressed genes were identified using DESeq2.^55^ Genes were considered significantly upregulated if they exhibited a log_2_ fold-change ≥ 1 and a false discovery rate (FDR) < 0.05, and significantly downregulated if they exhibited a log_2_ fold-change ≤ -1 and FDR < 0.05.

#### Cell viability assay

Cell viability was assessed using the Cell Counting Kit-8 (ApexBio Technology, K1018) following the manufacturer’s protocol. Briefly, 2×10^3^ cells were seeded onto a 96-well plate. At each time point, 10 μL of CCK-8 reagent was added to wells containing 100 μL of cell culture medium, and the mixture was incubated for 2 hours at 37°C in a humidified incubator. Absorbance at 450 nm was recorded using the Infinite M200 Plex microplate reader (Tecan) as a surrogate for cell viability.

#### Subcutaneous tumor implantation

Single-cell suspensions were harvested by trypsinization, washed with PBS, and resuspended in DPBS (Corning, 21031CV) with 1 g/L glucose to achieve a final concentration of 3×10^6^ cells/mL. A total of 50 μL of cells per mouse was injected subcutaneously into the right flank of an 8-week-old female mouse, and tumor size was monitored with a caliper every 3 days. Mice were euthanized by CO_2_ inhalation once the tumor reached 2 cm in any dimension according to IACUC instructions. The tumor size was calculated using the formula: Volume = (π/6) × Length × Width × Height.

#### Orthotopic tumor implantation

Single-cell suspensions were collected as above to achieve a final concentration of 2.5×10^6^ cells/mL. A total of 20 μL of cell suspension per mouse was injected orthotopically into the pancreas of female mice aged 6–9 weeks. Mice were euthanized by CO_2_ inhalation at the same endpoint for tumor size assessment. For survival analysis, mice were euthanized when signs of distress were observed, and the time to death or euthanasia was used as the survival endpoint.

#### Sample and library preparation for single-cell RNA sequencing (scRNAseq)

Tumor-bearing mice were sacrificed on day 15 after cell inoculation. Pancreatic tumors with spleens from 12 mice were dissected, weighed, and rinsed. To avoid cell contamination from the normal pancreas and spleen, half of the tumor from each mouse was isolated from the opposite side of the spleen. To reduce individual variance among mice, two bisected tumors within each group were randomly combined. A total of 3 samples for each group were dissociated to prepare single-cell suspensions.

Tissue digestion is identical to the method described in the tissue dissociation section above. To achieve greater than 85% cell viability, we used a dead-cell removal kit (Miltenyi, 130-090-101) to remove dead cells and assess cell viability with trypan blue staining.

scRNAseq library preparation was performed according to the manufacturer’s guidelines (10x Genomics, user guide #CG000315 Rev E). Roughly 1×10^4^ cells were loaded into the microfluidic system to generate a gel bead-in-emulsion. More than 2×10^4^ read pairs per cell were captured. Sequencer-generated BCL files were converted to fastq format and subsequently aligned to the mm10 mouse genome using Cell Ranger (v7.2.0, 10x Genomics). Filtered output from Cell Ranger served as the basis for downstream analytical procedures.

#### Data processing of scRNAseq

The count matrices produced by Cell Ranger were loaded using the Read10X function from the Seurat package (v5.0.1)^56^ and subsequently converted into Seurat objects. Quality control filtering was based on gene detection thresholds and mitochondrial content: cells with fewer than 300 or more than 7,000 detected genes, or cell exhibiting mitochondrial UMI counts exceeding 10%, were excluded. Genes detected in fewer than three cells were also removed. Potential doublets were identified and filtered out using DoubletFinder (v2.0.3)^57^ with default parameters. Data normalization employed the NormalizeData function to standardize total expression across cells, followed by identification of the top 2,000 highly variable genes using TheFindVariableFeatures. The Seurat objects were scaled with ScaleData, incorporating regression of cell cycle effects based on phase scores calculated from known markers.^58^

Dimensionality reduction was performed using the RunPCA function to conduct principal component analysis. Subsequently, a shared nearest neighbor (SNN) graph was built based on selected principal components using the FindNeighbors and FindClusters functions from the Seurat package. Cell clustering was performed using a graph-based approach with the Louvain algorithm. Differentially expressed marker genes for each cluster were identified using the FindAllMarkers and FindMarkers functions.

Cell identity for each cluster was determined using a two-step approach that integrated marker-based and reference-based annotations. Initially, major cell types were manually assigned based on well-characterized canonical markers, including cancer cell markers (*Kras*, *Myc*, *Pvt1*, *Krt18*), immune cell markers (*Ptprc*), fibroblast markers (*Col1a1*, *Col1a2*), and endothelial cell markers (*Pecam1*, *Emcn*). *Ptprc*-positive cells were further annotated using the SingleR package^24^ with a reference dataset (main label). Identities assigned by SingleR were subsequently validated through canonical markers for macrophages (*Adgre1*, *Arg1*, *Itgax*), granulocytes (*S100a9*, *S100a8*, *G0s2*), monocytes (*Itgam*, *Cd14*), B cells (*Cd19*, *Cd79b*), NK cells (*Nkg7*), and T cells (*Cd3e*, *Cd3d*). Visualization was performed using the scCustomize package.^59^ Subclustering followed comparable procedures, including normalization, variable gene detection, dimensionality reduction, and clustering. Cancer-associated fibroblast (CAF) subsets were annotated based on canonical markers distinguishing myCAFs (*Postn*, *Lrrc15*, *Mmp11*, *Acta2*, *Thy1*), mesothelial cells (*Msln*, *Upk3b*, *Krt19*, *Lrrn4*), iCAFs (*Dpt*, *Pi16*, *Il6*, *Cxcl12*, *Cxcl1*), proliferating CAFs (*Tpx2*, *Ccna2*, *Cenpf*, *Cdc20*, *Mki67*), and antigen-presenting CAFs (*H2*-*Aa*, *H2*-*Ab1*, *Cd74*).

#### Immunohistochemistry and analysis

All reagents used in immunohistochemistry were purchased from VECTASTAIN Elite ABC HRP Kit (Vector Laboratories, PK-6101) unless otherwise stated. Tumor samples were harvested after mice were euthanized by CO_2_ inhalation and fixed with 4% paraformaldehyde in 0.1 M phosphate buffer (FD NeuroTechnologies, PF101) at 4°C for 24 hours and maintained in 70% ethanol solution at 4°C. Tissue sectioning was performed by the Tissue Management Shared Resource at the University of Texas Southwestern Medical Center. Sections of formalin-fixed, paraffin-embedded tissue were deparaffinized with Formula 83 (CBG Biotech, CH0104A) and rehydrated with graded alcohol. Antigen retrieval was performed using the Antigen Unmasking Solution (Vector Laboratories, H3300250) in a pressure cooker under 7.5 psi for 17 minutes. Slides were cooled down to room temperature, rinsed with Milli-Q water, and outlined with a PAP pen. Slides were blocked with BLOXALL Blocking Solution (Vector Laboratories, SP-6000) at room temperature for 10 minutes and washed with PBS. Slides were blocked with 1.5% normal goat serum in PBS at room temperature for 1 hour. The following primary antibodies were applied overnight at 4 °C: anti-POSTN (1:2,000, Cell Signaling Technology, 35126S), anti-CD3E (1:500, Thermo Fisher Scientific, MA5-35204), and anti-CD8A (1:500, Cell Signaling Technology, 98941S). Slides were washed with PBST and incubated with biotinylated secondary antibody, followed by sequential application of ABC Reagent. The DAB Substrate Kit (Vector Laboratories, SK-4105) was used for visualization. Finally, slides were counterstained with hematoxylin (Vector Laboratories, H-3401-500) and mounted with mounting medium (Vector Laboratories, H-5501-60). After drying at room temperature for 24 hours, slides were scanned by the Nanozoomer S60. To quantify DAB staining intensity, four images (each 0.715 mm²) were acquired from different regions of each sample using NDP viewer (v2.9.29). The staining intensity was defined as the ratio of DAB-positive area to the total tissue area. Image processing was performed using CellProfiler (v4.2.8). Briefly, original colored images were converted to grayscale, and the threshold was adjusted to segment DAB-positive regions and blank areas. Blank areas were excluded by subtracting non-tissue regions from the total image area.

#### Immunofluorescence on FFPE sections

Slides were treated in the same way as in immunohistochemistry experiments, from deparaffinization to antigen retrieval. Then, samples were blocked with 1.5% normal goat serum in PBS. The following antibodies were incubated overnight at 4°C: anti-RAS (1:200, Cell Signaling Technology, 91054), anti-MYC (1:100, Abcam, ab32072), anti-POSTN (1:2000, Cell Signaling Technology, 35126S), and anti-KI67 (1:100, Thermo Fisher Scientific, 14-5698-80). After washing with PBST, the following secondary antibodies were applied at room temperature for 1 hour: anti-Rabbit antibody conjugated with AlexaFluor 594 (1:1,000, Thermo Fisher Scientific, A32740), anti-Rabbit antibody conjugated with AlexaFluor 488 (1:1,000, Thermo Fisher Scientific, A32731), anti-Mouse antibody conjugated with AlexaFluor 594 (1:1,000, Thermo Fisher Scientific, A32742), and anti-Rat antibody conjugated with AlexaFluor 568 (1:1,000, Thermo Fisher Scientific, A11077). After washing with PBST buffer, autofluorescence was quenched using TrueVIEW reagents (Vector Laboratories, SP-8400-15). Sections were counterstained with DAPI and mounted with antifade medium (Thermo Fisher Scientific, P36961). The MYC protein fluorescence intensity was quantified using CellProfiler (v4.2.8). Briefly, nuclei were identified based on the DAPI channel, and the resulting nuclear masks were applied to the MYC protein fluorescence channels. The fluorescence intensity of the MYC signal within each nucleus was measured. The RAS protein fluorescence intensity was quantified in ImageJ by manually outlining each cell and measuring the fluorescence intensity in the RAS signal channel.

#### OligoPaint DNA FISH on FFPE sections

DNA FISH on formalin-fixed paraffin-embedded (FFPE) tissue sections was carried out following the steps outlined in our previously published protocol.^60^ Briefly, FFPE sections were deparaffinized with Formula 83 and rehydrated with an ethanol series. Tissues were then treated with 0.2N HCl at room temperature for 20 minutes, 10 mM citric acid at 90°C for 20 minutes, and proteinase K at room temperature for 1 minute. After dehydration with an ethanol series, OligoPaint FISH was performed as described above. Finally, autofluorescence was eliminated using the TrueVIEW Autofluorescence Quenching Kit (Vector Laboratories, SP-8400) before DAPI counterstaining. The FISH signal intensity was quantified using CellProfiler (v4.2.8). Briefly, nuclei were identified based on the DAPI channel, and the resulting nuclear masks were applied to the FISH signal channels. The FISH signals were refined into sharp puncta by threshold adjustment, and the total signal intensity within each nucleus was measured.

To combine OligoPaint DNA FISH and IF in the same section, OligoPaint DNA FISH was first performed as described above. After secondary probe hybridization and washing, slides were blocked with 10% normal goat serum with 0.05% Triton X-100 in PBS for 1 hour, followed by immunostaining as described in the immunofluorescence section.

#### Fluorescence microscopy

DNA FISH and immunofluorescence imaging were captured using a Zeiss Axio Observer 7 microscope equipped with the Apotome 3 optical sectioning system. Images were acquired with a 63× Plan-Apochromat oil immersion objective lens (NA 1.40) and processed using Zeiss ZEN software (version 3.4). For each sample, at least five z-stack images were captured at 1 μm intervals, followed by the generation of maximum intensity projections to produce 2D representations. Whole tissue scans of FISH or combined FISH and immunofluorescence staining were performed using a Zeiss Axioscan 7 slide scanner with a 40× objective.

#### Gene signature, immune infiltration analysis, and survival analysis

*Kras* super-expressor signature genes (Supplementary Table 1) were identified using Seurat (v5) based on the following criteria: log_2_ fold-change ≥ 1.5, presence of human orthologs, and ranking among the top 30 genes by FDR. These genes were mapped to their human orthologs to retrieve transcript per million (TPM) values from the TCGA-PAAD cohort, obtained via UCSC Xena. *Kras* super-expressor signature scores were calculated as the geometric mean of TPM values for the corresponding human orthologs. Similarly, myCAF scores were defined as the geometric mean of TPM values for a curated list of myCAF biomarker genes (Supplementary Table 2).

Immune infiltration scores were sourced from TIMEDB. Pearson correlation analysis was performed to assess association significance, and p-values were adjusted using the FDR method to control for false positives.

Overall survival analysis was performed using GEPIA2 (gepia2.cancer-pku.cn).^61^ The cutoff for distinguishing low and high expression was set as 50% of the median.

#### RNAi

*Kras* siRNA (Thermo Fisher Scientific, s68936) was delivered to cells using Lipofectamine RNAiMAX (Thermo Fisher Scientific, 13778150) according to the manufacturer’s protocol. Transfected cells were harvested 48 hours after transfection for subsequent experimental analyses.

#### Western blot

Cells treated with inhibitors were harvested and lysed in RIPA buffer (Boston BioProducts, BP-115) with protease inhibitor (Roche, 04693116001) and phosphatase inhibitor (Roche, 04906837001). Protein concentration was determined using the BCA Protein Assay Kit (Thermo Fisher Scientific, A53225). Equal amount of protein samples were prepared in 1× Laemmli sample buffer (Bio-Rad, 1610747) and heated at 95°C for 10 minutes, ran on 4-20% Mini-PROTEAN gels (Bio-Rad, 4568096), and transferred to a nitrocellulose membrane using Trans-Blot Turbo system with transfer kit (Bio-Rad, 1704270). The membrane was blocked with 5% BSA in TBS with 0.1% Tween-20 (Fisher Scientific, BP337-500) for 1 hour. The following primary antibodies were used: anti-pMAPK (1:2,000, Cell Signaling Technology, 4370T), anti-MAPK (1:1000, Cell Signaling Technology, 4695T), anti-p4E-BP1 (1:1,000, Cell Signaling Technology, 2855T), anti-4E-BP1 (1:1,000, Cell Signaling Technology, 9644T), and anti-Vinculin (1:10,000, Proteintech, 66305-1-Ig). The membranes were washed 3 times with TBST and incubated with HRP-conjugated secondary antibodies (Anti-mouse, 1:5,000, Cell Signaling Technology, 7076S; Anti-rabbit, 1:5,000, Cell Signaling Technology, 7074S) for 1 hour. Membranes were washed 3 times with TBST, then visualized with a chemiluminescent substrate (Thermo Fisher Scientific, 34580) and imaged using ImageQuant 800 (Amersham).

#### RT-qPCR

RNA was extracted using the Quick-RNA Miniprep Plus Kit (Zymo Research, 50-125-1683) and quantified by NanoDrop (Thermo Scientific). One microgram of RNA was used for reverse transcription to generate cDNA using the Maxima H Minus reverse transcriptase (Thermo Scientific, EP0753). Primers used are in the table of the method section. The cDNA was added with 1× SYBR Green qPCR Master Mix (Selleck Chemicals, B21203) and 400 nM of the respective forward and reverse primers for qPCR on a CFX Opus 96 System (Bio-Rad). The relative RNA level was calculated by the ddCt method.

#### CRISPR knockout

Two sgRNAs were designed using E-CRISP,^62^ synthesized by Integrated DNA Technologies, and cloned into the LentiCRISPRv2 vector (Addgene, 52961). Lentivirus packaging was performed by co-transfecting HEK293T cells with lentiviral transfer plasmids and helper plasmids (pMD2.G, Addgene, 12259; psPAX2, Addgene, 12260). The CT1BA5EC cells were infected with the lentivirus and selected by puromycin (InvivoGen, ant-pr-1). Single-cell clones were isolated using a flow cytometer (BD FACSAria Fusion) and genotyped to confirm homozygous knockout.

#### Xenium *in situ* high-plex assay

We developed a fully customized 480-gene panel (Supplementary Table 3) for cell type identification and functional characterization. FFPE tissue blocks were sectioned at 5-μm thickness and mounted within the designated capture area (10.45 mm × 22.45 mm) of Xenium slides (10x Genomics). Sample preparation followed the manufacturer’s protocol for FFPE tissues (10x Genomics Xenium In Situ FFPE Tissue Preparation Guide). Mounted sections underwent deparaffinization in xylene followed by gradient rehydration through graded ethanol washes. Enzymatic permeabilization was then performed to enhance mRNA accessibility.

Target mRNA hybridization was conducted overnight at 50°C using gene-specific probes, after which excess unbound probes were washed away. Locked nucleic acid probes annealed to target transcripts were subsequently ligated using Xenium ligase A/B at 37°C for 2 hours. Gene-specific circular probes were then amplified by rolling-circle amplification to generate multiple copies of unique molecular barcodes. Following additional wash steps, autofluorescence was eliminated through chemical quenching. Imaging was performed on a Xenium analyzer (10x Genomics), with DAPI counterstaining providing nuclear morphology for cellular boundaries. Cell segmentation was accomplished using the instrument’s integrated machine learning algorithms, and the resulting data files were processed for subsequent computational analysis.

#### Xenium data processing and analysis

Xenium spatial transcriptomics data were analyzed using Seurat (v5.3.0) following standard workflows with modifications. Cell-by-gene expression matrices generated by the Xenium analyzer were imported into R and consolidated into a unified Seurat object. Quality control filtering was applied to remove low-quality cells, followed by variance-stabilizing transformation and normalization via the SCTransform method.

For unsupervised cell type identification, we performed dimensionality reduction via principal component analysis on highly variable genes (RunPCA). A k-nearest neighbor graph was constructed in the reduced-dimensional PCA space (FindNeighbors), followed by graph-based clustering using the Louvain algorithm (FindClusters, resolution = 0.2). Cell type identities were assigned to each cluster based on the expression of canonical marker genes. Clustering results were visualized in both reduced-dimensional space via Uniform Manifold Approximation and Projection (UMAP) and in their original spatial coordinates to preserve tissue architecture.

#### Statistics

All experiments were performed for at least three times. Statistical analyses were detailed in the corresponding figure legends. For data that were normally distributed and exhibited homoscedasticity, Student’s t-test or ANOVA was used. Otherwise, the corresponding statistical analysis was indicated in the figure legend. P values derived from multiple comparisons were adjusted using the false discovery rate (FDR) method. All tests were two-sided. Box plots display the median and interquartile range (IQR), with whiskers extending to 1.5× the IQR. Data points and error bars shown on line plots represent the mean ± standard error of the mean (SEM).

#### Data availability

All sequencing data will be uploaded to the NIH Sequence Read Archive repository and made publicly accessible upon the acceptance of this manuscript.

## Author contributions

**Kailiang Qiao:** Investigation, Validation, Formal analysis, Visualization, Writing - Review & Editing. **Qing-Lin Yang:** Investigation, Validation, Formal analysis, Visualization, Writing - Review & Editing. **Tuo Li:** Investigation, Writing - Review & Editing. **Xiongfeng Chen:** Formal analysis. **Zeynep Yazgan:** Investigation. **Yoon Jung Kim:** Investigation. **Collin Gilbreath:** Investigation. **Jun Yi Stanley Lim:** Investigation. **Yipeng Xie:** Investigation. **Xiaohui Sun:** Investigation. **Yang Liu:** Investigation. **Yiyue Jia:** Investigation. **Zhijian J. Chen:** Conceptualization, Writing - Original Draft, Funding acquisition. **Huocong Huang:** Conceptualization, Methodology, Formal analysis, Writing - Original Draft, Funding acquisition. **Sihan Wu:** Conceptualization, Methodology, Resources, Formal analysis, Writing - Original Draft, Funding acquisition, Visualization, Supervision.

## Acknowledgement

Sihan Wu is supported by the Cancer Prevention and Research Institute of Texas (CPRIT, RR210034) and the American Cancer Society (CAT-24-1379043-01-CAT). Huocong Huang is supported by the National Cancer Institute (R00 CA252009). Jun Yi Stanley Lim is supported by the CPRIT Training Grant (RP210041). This work was delivered as part of the eDyNAmiC team supported by the Cancer Grand Challenges partnership funded by Cancer Research UK (P.S.M CGCATF-2021/100012, Z.J.C., S.W. CGCATF-2021/100023) and the National Cancer Institute (P.S.M. OT2CA278688, Z.J.C., S.W. OT2CA278683). Work in the Chen laboratory is supported by grants from the National Cancer Institute (R01CA299257) and the Welch Foundation (I-1389). We acknowledge the assistance of the University of Texas Southwestern Tissue Management Shared Resource, a shared resource at the Simmons Comprehensive Cancer Center, which is supported in part by the National Cancer Institute under award number P30 CA142543. We acknowledge the assistance of the University of Texas Southwestern Whole Brain Microscopy Facility, RRID: SCR_017949. Certain graphic elements were obtained from the open-sourced Bioicons and NIH BIOART (CC-BY 3.0 unported license).

## Supplementary Figure Legends

**Figure S1. Amplicon structure of EC and HSR PDAC cell lines.**

The structures of *Kras* and *Myc* amplicons in EC1/2 and HSR1/2 PDAC cell lines were analyzed using AmpliconArchitect, showing high similarity between ecDNA and HSR amplicons.

**Figure S2. Molecular characteristics of EC and HSR PDAC cell lines.**

(A) Stacked bar plot showing the percentage of *KRAS^G12D^* allele in EC1/2 and HSR1/2 isogenic pairs.

(B) Principal component analysis (PCA) of bulk-cell gene expression profiles showing clear separation between EC and HSR cell lines. Rep1 and Rep2 indicate two sequencing replicates.

(C) Heatmap showing that only 1% of genes in the whole transcriptome were differentially expressed among EC and HSR groups.

(D) GSEA analysis indicating minimal differences for KRAS and MYC signaling between the EC and HSR transcriptome in bulk-cell cultured *in vitro*.

(E) Cell viability assay *in vitro* using CCK-8. Data represent mean ± SEM from three independent experiments. Two-way ANOVA test with Tukey’s HSD. NS indicates not significant.

**Figure S3. Expression of canonical biomarkers of each cell population**

(A) Gross images and tumor weight quantification of tumors for scRNAseq. The mid-stage tumor weight between EC1 and HSR1 has not shown significant differences. Student’s t-test.

(B) Dot plot showing expression of canonical marker genes across distinct cell populations to validate the cell type annotation identified by SingleR. Dot size indicates the percentage of cells expressing the gene within each cell type, and dot color represents the scaled expression level.

(C) Dot plot showing expression of canonical marker genes across CAF subtypes to annotate the subclusters after dimension reduction. Dot size indicates the percentage of cells expressing the gene within each cell type, and dot color represents the scaled expression level.

(D) UMAP visualization of T cell subclusters.

(E) Dot plot showing expression of canonical T cell markers.

(F) Cell composition analysis of T cell subtypes in EC and HSR groups. Fisher’s exact test.

(G) Representative images of CD3E IHC staining for mid-stage tumors showing both low- and high-magnification views. Red letter S and circles indicate the spleen tissue areas as a positive control for CD3E staining. Scale bars 5 mm (low magnification), 50 μm (high magnification).

**Figure S4. Variance and correlation analyses among ecDNA copy number, mRNA transcription, and protein expression.**

(A) Representative *Kras* DNA FISH images taken from FFPE tumor sections. Scale bar: 10 μm.

(B) Density plot showing the distribution of *Kras* DNA FISH signal intensity of FFPE tumor sections in each cell. The variances of EC and HSR were assessed using MAD and IQR. Number of nuclei analyzed: EC, n = 984; HSR, n = 776.

(C) Representative RAS protein IF images taken from FFPE tumor sections. Scale bar: 10 μm.

(D) Density plot showing the distribution of RAS protein IF intensity of tumor section in each cell. The variances of EC and HSR were assessed using MAD and IQR. Number of cells analyzed: EC, n = 288; HSR, n = 402.

(E) Representative images of RAS protein IF and *Kras* DNA FISH co-staining in FFPE tumor sections. Scale bar: 10 μm.

(F) Correlation analysis between the intensity of RAS protein IF and the intensity of *Kras* DNA FISH in FFPE tumor sections. Spearman correlation test. Number of cells analyzed: n = 723.

(G) Density plot of *Myc* mRNA expression of EC and HSR cells from scRNAseq data. The distribution variance was assessed using MAD and IQR. Number of cells analyzed: EC, n = 10,000; HSR, n = 10,000.

(H) Representative images of MYC protein IF and *Myc* DNA FISH co-staining in FFPE tumor sections. Scale bar: 10 μm.

(I) Correlation analysis between the intensity of MYC protein IF and the intensity of *Myc* DNA FISH in FFPE tumor sections. Spearman correlation test. Number of cells analyzed: n = 977.

**Figure S5. Comparative analysis of *Kras* super-expressors and normal-expressors.**

(A) Schematic workflow for comparative analysis of *Kras* super-expressors and normal-expressors.

(B) Density plot of *Kras* mRNA expression from 10,000 randomly selected cancer cells, each from EC and HSR tumors. Cells with the top 5% *Kras* expression were defined as *Kras* super-expressors. GSEA analysis showed that KRAS signaling was significantly upregulated in the super-expressor compared with the normal-expressor, but not between EC and HSR in the normal-expressor.

(C) UMAP view displaying the *Kras* super-expressor cells in the EC and HSR groups.

(D) Dot plot showing the expression of selected replication stress marker genes between *Kras* super-expressors and normal-expressors.

(E) Correlation analysis between the intensity of *Kras* DNA FISH and the intensity of *Myc* DNA FISH in FFPE EC tumor sections. Spearman correlation test. Number of nuclei analyzed: n = 1284.

(F) Schematic illustrating the inheritance patterns of ecDNA and chromosomes. Asymmetric mitotic segregation of ecDNA can rapidly generate oncogene super-expressors, promoting tumor growth. However, elevated oncogene dosage induces cellular stress and reduces fitness. The dynamic nature of ecDNA inheritance enables reversible transitions between super- and normal-expressors, enhancing cancer cell adaptability to the tumor microenvironment. In contrast, the stable inheritance of chromosomes restricts the emergence of super-expressors and limits their interconversion with normal-expressors.

**Figure S6. Effector gene analysis in *Kras* super-expressors.**

(A) Dot plot showing the differential expressions of *CEACAM1*, *PDCD1LG2*, *PTHLH,* and *CSF2* between TCGA-PAAD tumor samples and GTEx normal tissues. Wilcoxon test.

(B) Overall survival of TCGA-PAAD cohort stratified by *CEACAM1*, *PDCD1LG2*, *PTHLH*, and *CSF2* expression (median 50% cutoff). Log-rank test.

(C-D) Violin plot showing the Log1p normalized mRNA expression of *Areg* and *Csf2* in super- and normal- expressors. Wilcoxon test with FDR correction.

(E) Violin plot showing the Log1p normalized mRNA expression of *Areg* in EC and HSR cells. Wilcoxon test with FDR correction.

(F) qPCR detection of *Areg* and *Kras* mRNA levels in EC cells with or without transfection of *Kras* siRNA and siRNA control. Student’s t-test.

(G) Western plot showing the expression of MAPK and pMAPK (T202/Y204) in EC cells treated with trametinib and SCH772984 for 24 hours in a dose-dependent way.

(H) Western plot showing the expression of 4E-BP and p4E-BP (T37/46) in EC cells treated with Rapamycin and Torin 2 for 24 hours in a dose-dependent way.

(I) qPCR detection of *Areg* mRNA level in EC cells treated with vehicle control DMSO, SCH772984 (2 μM), Trametinib (40 nM), Rapamycin (0.125 μM), Torin2 (200 nM), and combination treatment with Trametinib (40 nM) and Torin2 (200 nM) for 24 hours. One-way ANOVA with Tukey’s HSD correction.

(J) Schematic image showing the KRAS signaling pathways.

**Figure S7. *Areg* knockout validation and extended data for Figure 4.**

(A) qPCR detection of *Areg* mRNA level in two *Areg-*knockout clones (sg*Areg*#1 and #2) compared to non-targeting sgRNA control (sgNTC).

(B) *Kras* and *Myc* DNA FISH on metaphase spreads from *Areg*-knockout cells. Scale bar: 5 μm.

(C-D) *Kras* and *Myc* ecDNA counts in sgNTC and sg*Areg* clones. One-way ANOVA revealed no significant difference. Sample sizes: sgNTC, n = 31; sg*Areg*#1, n = 35; sg*Areg*#2, n = 32.

(E) *In vitro* cell viability of sgNTC and sg*Areg* clones measured by CCK-8. Two-way ANOVA computed p-values with Tukey’s HSD correction.

(F) Log1p normalized *Areg* and *Egfr* mRNA expression among different cell populations detected by scRNAseq.

(G) Representative images of POSTN and KI67 protein IF staining. Blue arrows indicate the KI67-negative myCAFs, and yellow arrows indicate KI67-positive myCAFs. Scale bar: 50 μm (low magnification), 20 μm (high magnification).

(H) Percentage of the KI67-positive myCAFs among total myCAFs. One-way ANOVA test with Tukey’s HSD correction. More than 100 POSTN+ myCAFs per mouse were counted. The p-value indicates the significance compared sgNTC. Sample sizes: sgNTC, n = 7; sg*Areg*#1, n = 7; sg*Areg*#2, n = 7.

(I) Pearson correlation analysis between the myCAF score (calculated as the geometric mean of TPMs of myCAF biomarker genes) and immune cell infiltration scores obtained from TIMEDB, across samples in the TCGA-PAAD cohort.

**Figure S8. Spatial organization of cells with *Kras* amplification in autochthonous *KPfC* tumor.**

Representative images of the whole slide scanning stained by *Kras* DNA FISH from an autochthonous *KPfC* mouse tumor.

**Figure S9. Spatial organization of regions with high- and low-*KRAS* expression in human PDAC.**

(A) Density map of *KRAS* expression in the entire section from one human PDAC tumor (left) and cell identity analyzed by Xenium spatial transcriptomics (right). The locations of the selected regions for cell density analysis, shown in Figures 5C-D, were marked by red (*KRAS*-high) and yellow (*KRAS*-low) squares. Scale bar: 1 mm.

(B) Dot plot showing marker gene expression of major cell populations identified by Xenium spatial transcriptomics in a human PDAC sample.

(C) Subcellular spatial mapping of transcripts for *EPCAM*, *KRAS*, and *AREG* in a representative region with *KRAS*-high or *KRAS*-low cancer cells. Scale bar: 50 μm.

