## Supplementary figures and images for "ecDNA-driven oncogene super-expressors shape immunoevasive tumor microenvironment"

### supplementary figures

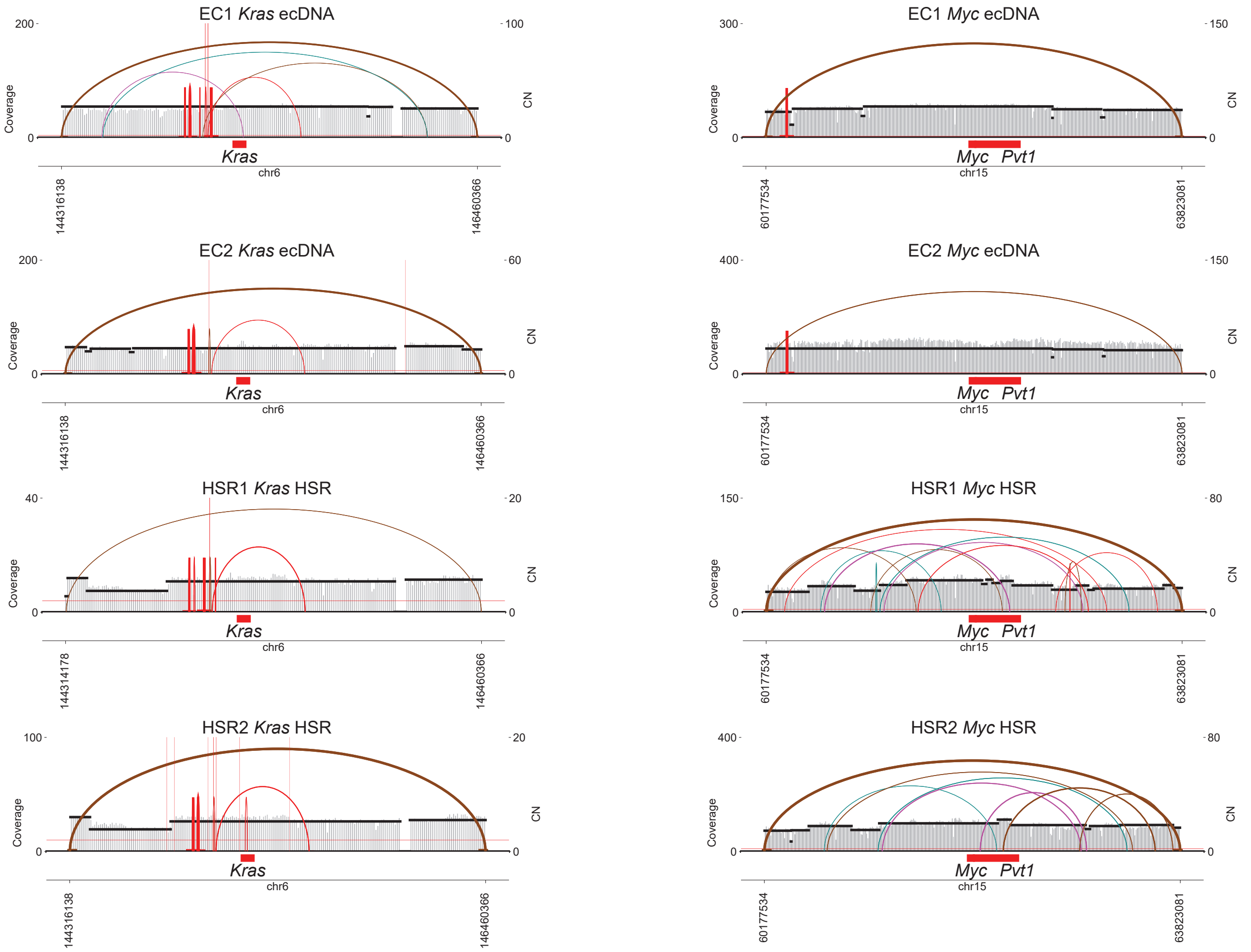

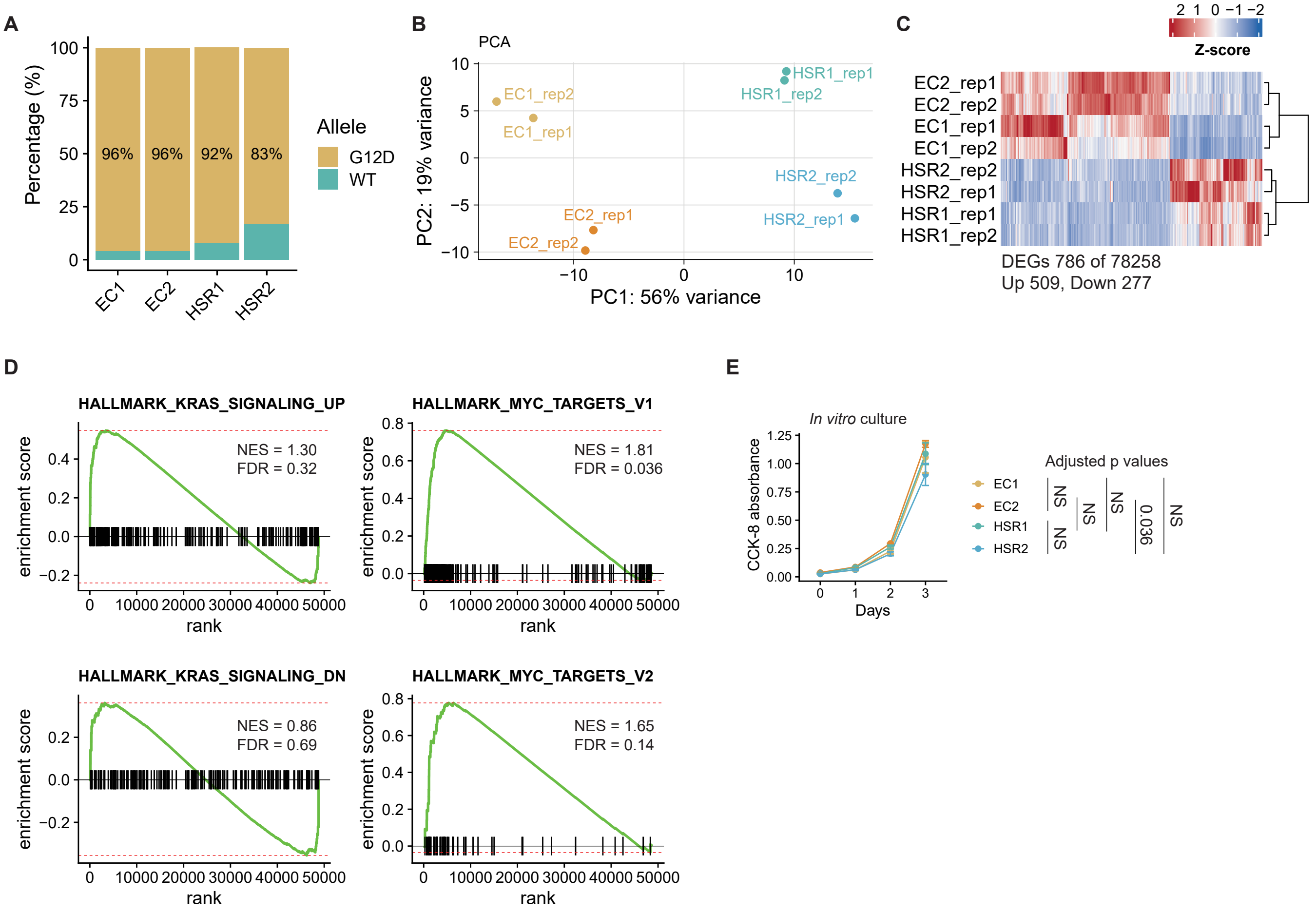

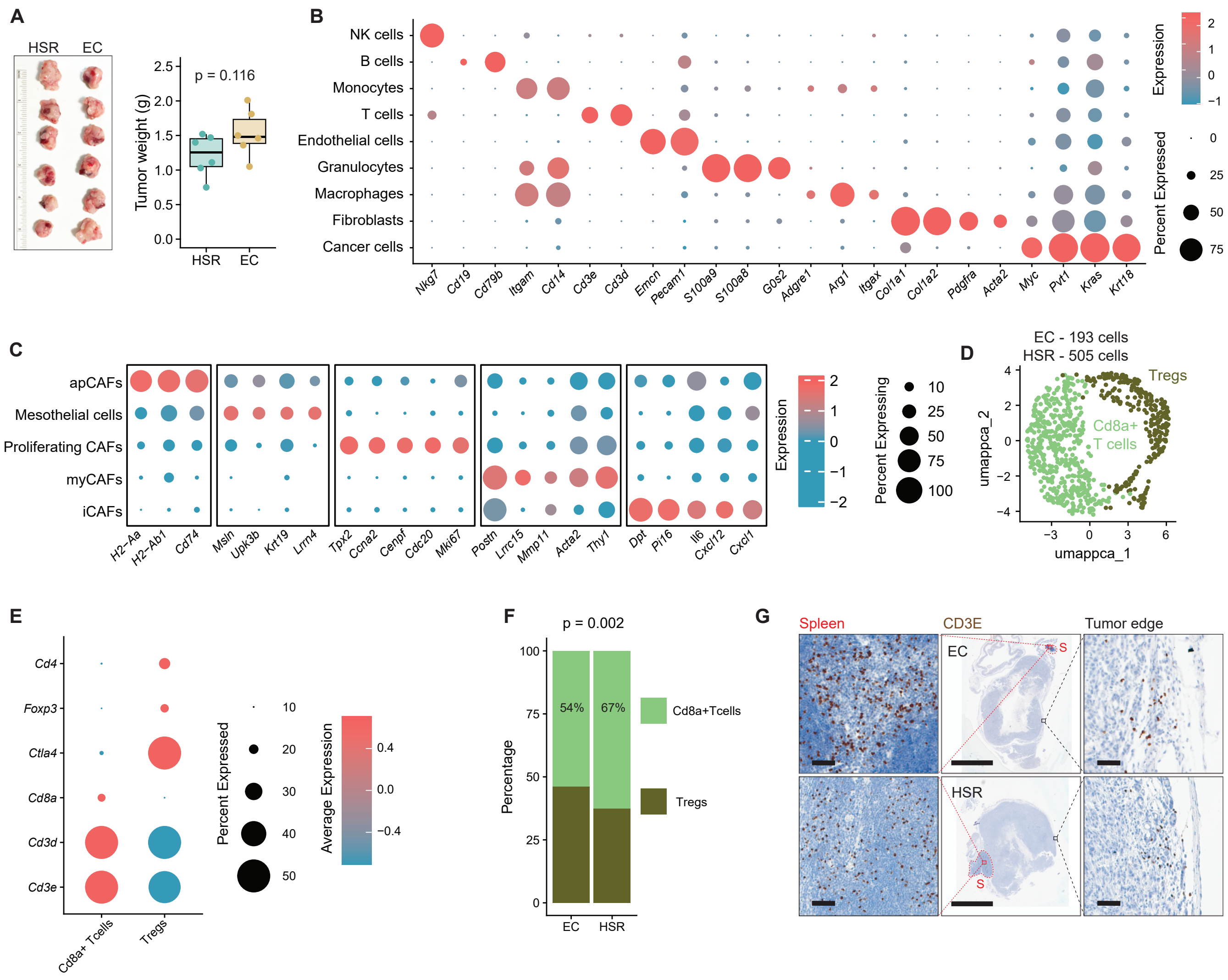

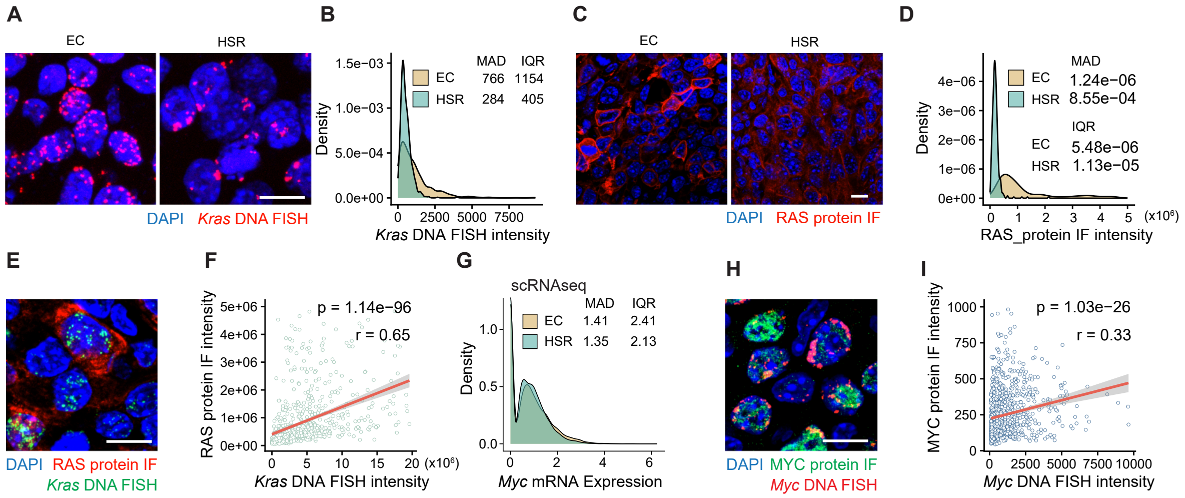

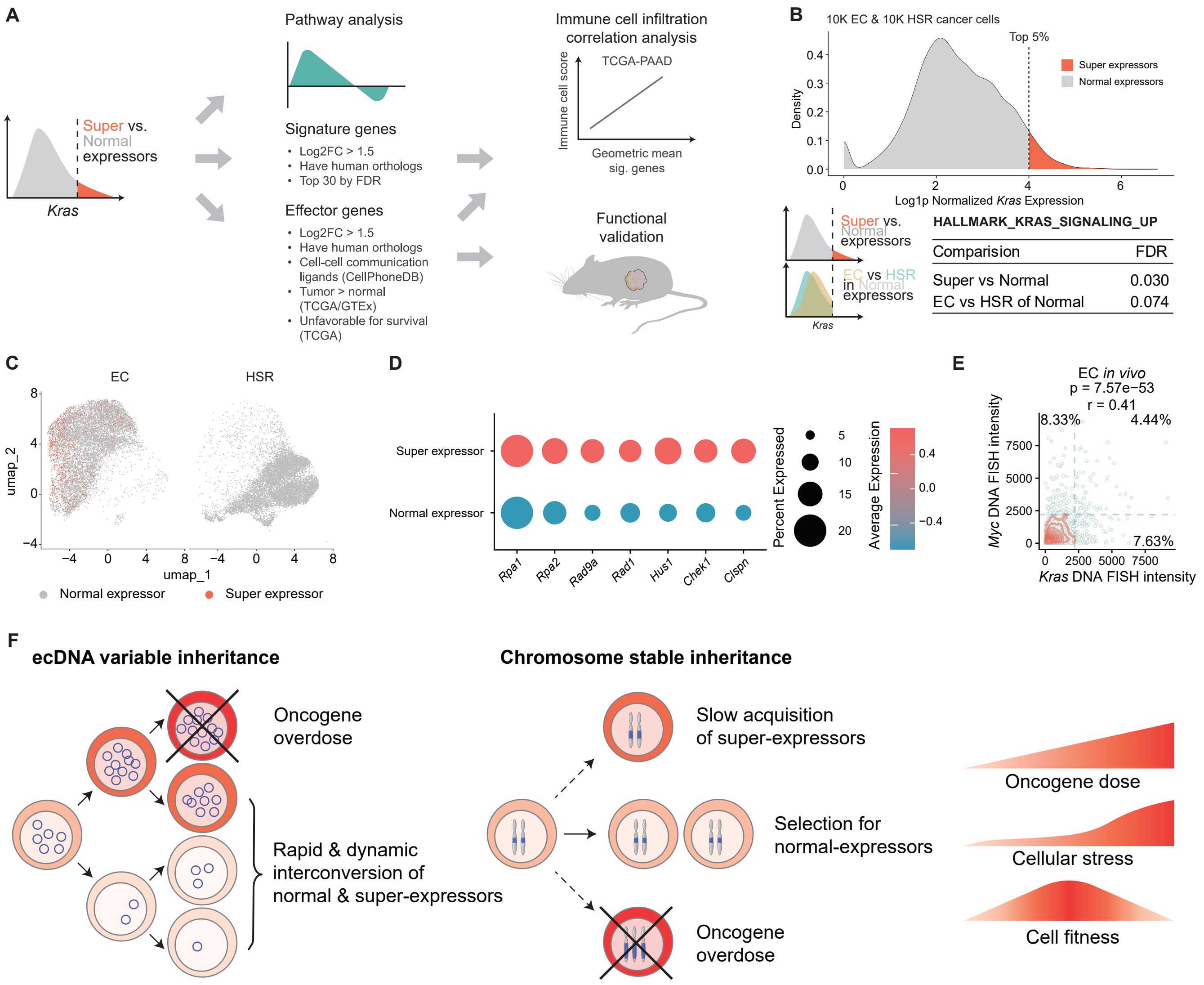

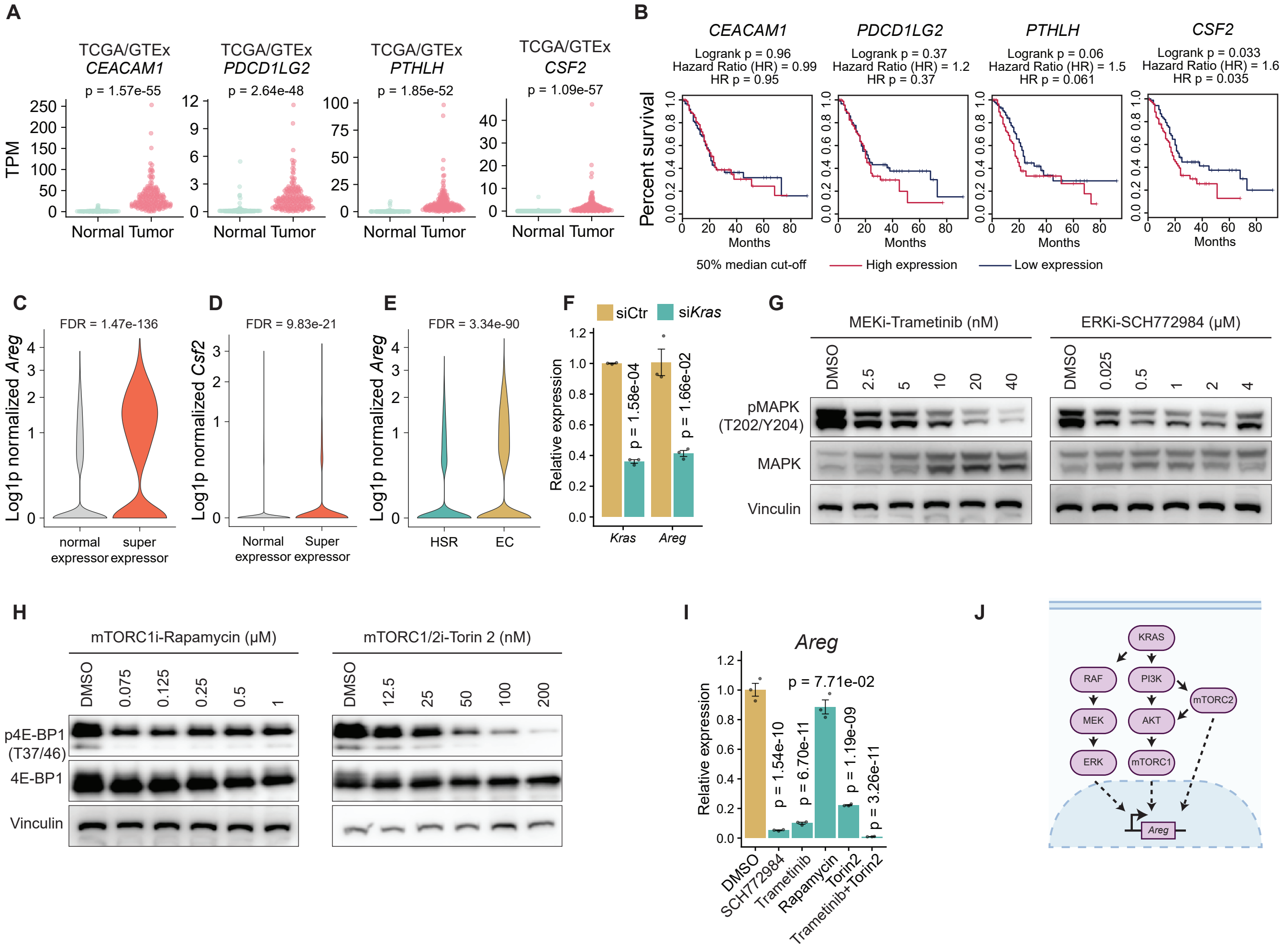

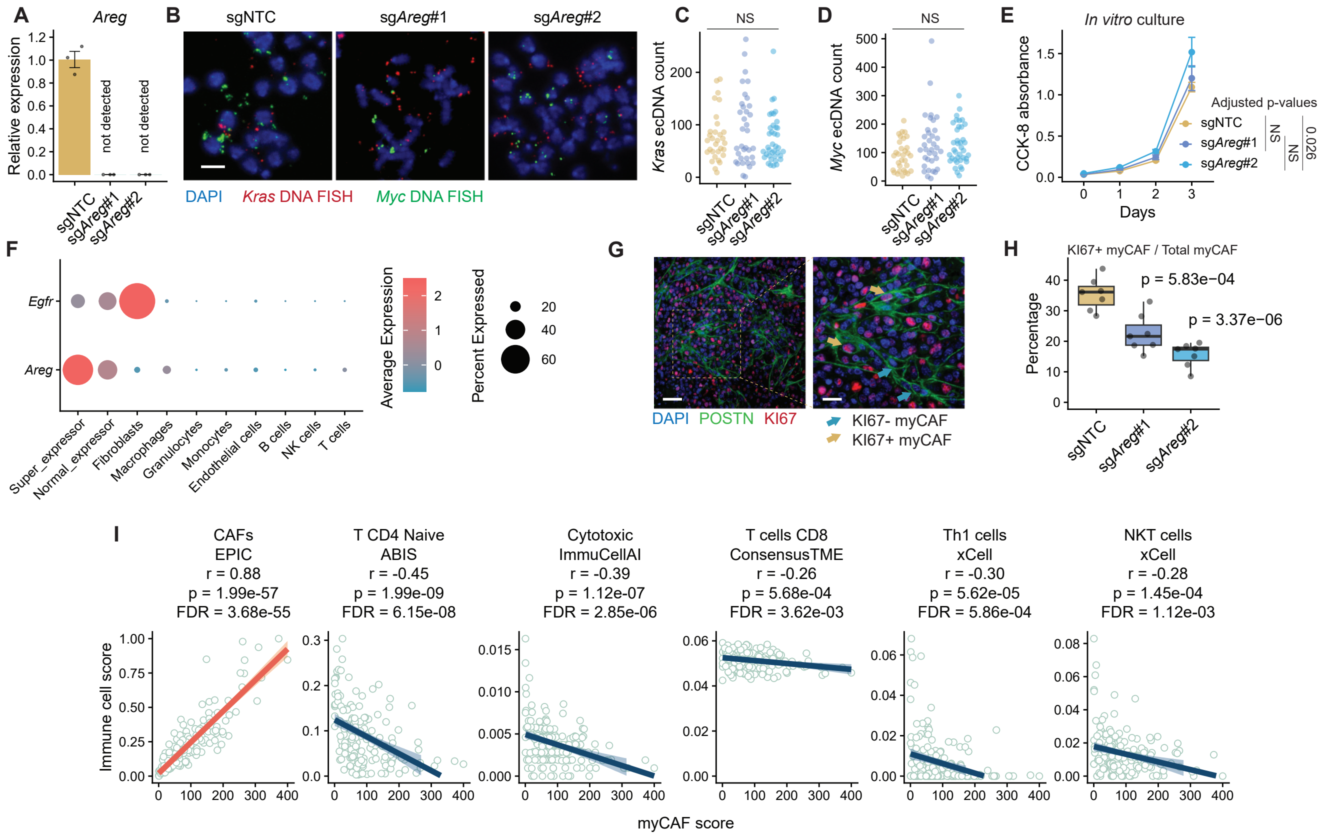

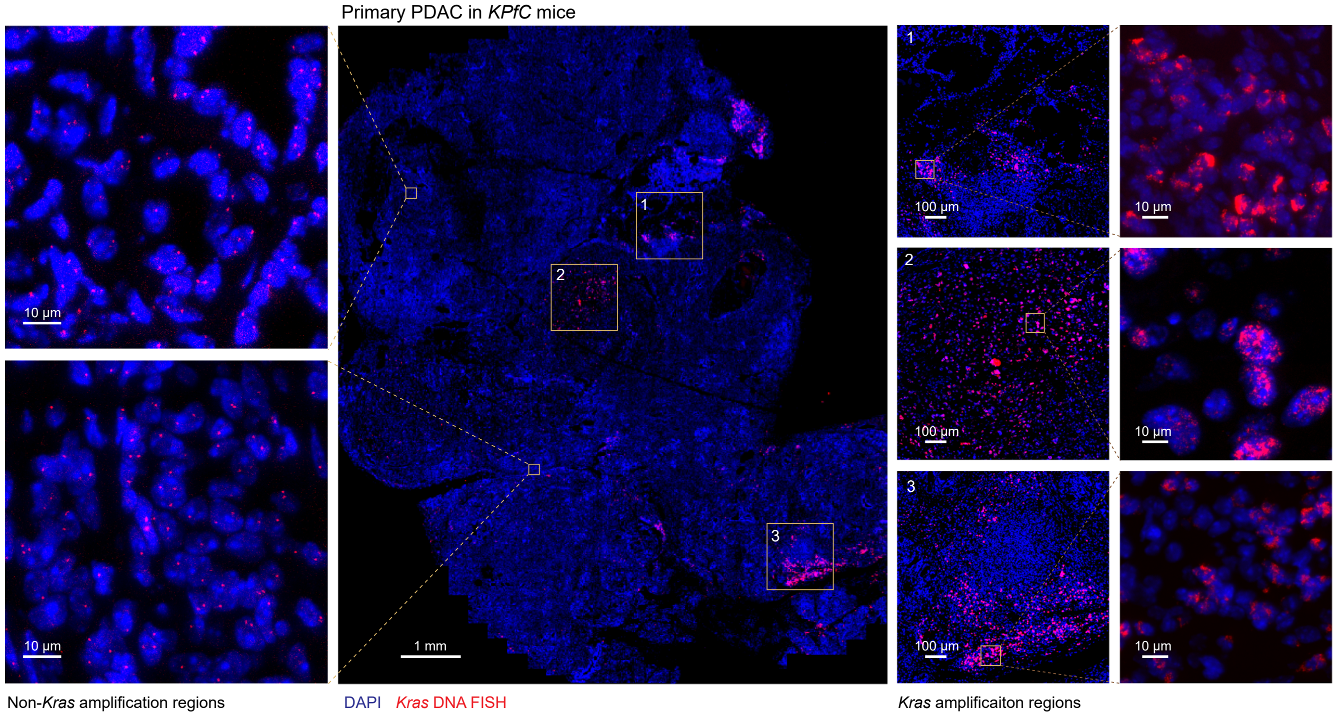

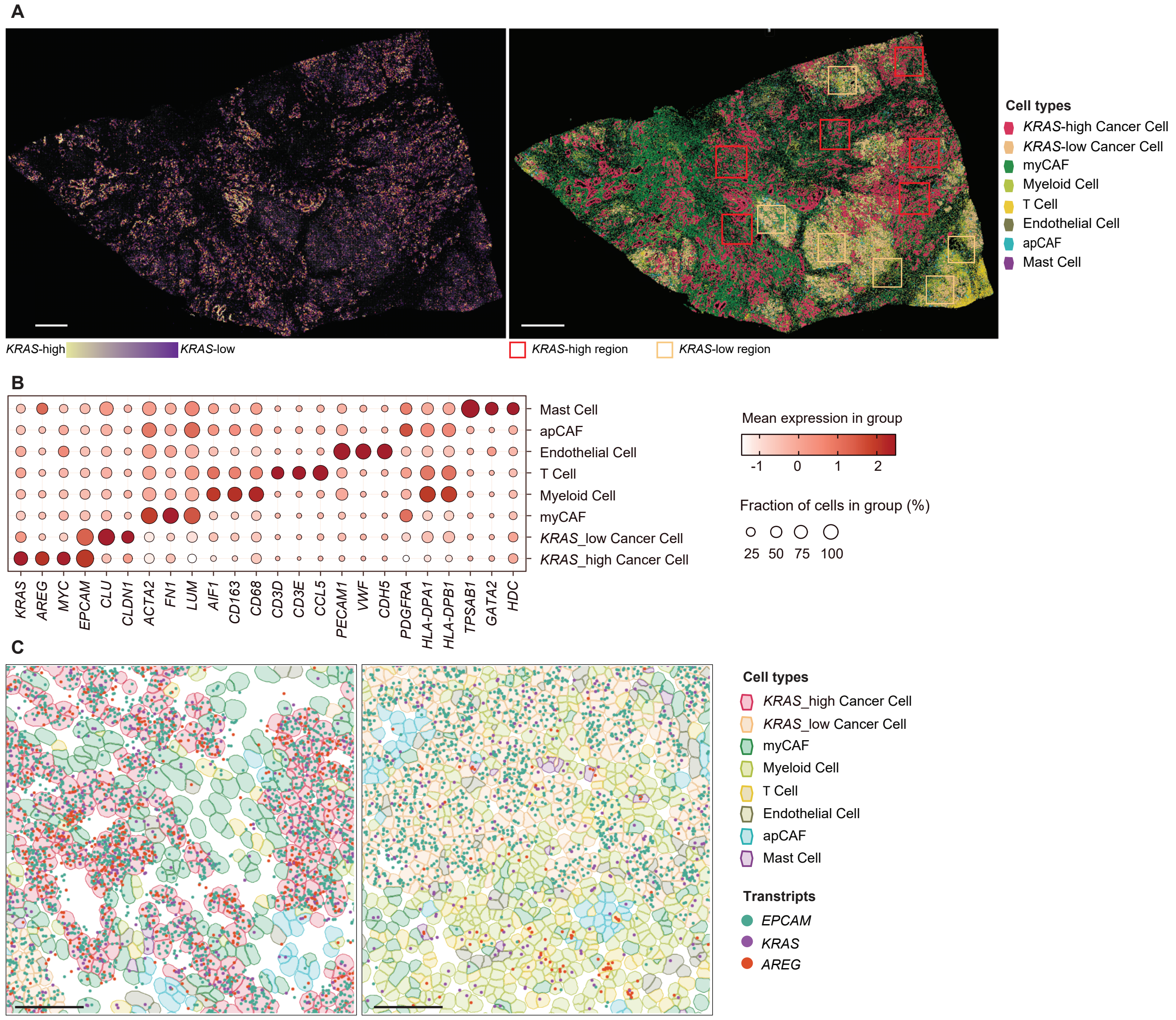
